# Sol-gel Transition Drives Hyper-fast Mixing in a Giant Cell

**DOI:** 10.64898/2026.07.13.738335

**Authors:** Ulises Diaz, Moumita Das, Sameer Thukral, Judy Abuel, Mikesha Carter, Alba Marino, Laura Galvan, Austin Irungu, Jasmine Leiva, Alexander Ballor, Wallace F. Marshall

## Abstract

The cytoplasm is a crowded and dynamic fluid within which cellular building blocks such as mRNA, proteins, or organelles undergo transport and mixing. Although small things like proteins can eventually mix through diffusion, the high viscosity of cytoplasm means that it should be difficult to obtain significant mixing for structures in the size range of mRNA, multi-protein complexes or organelles. In large amoeboid cells, the cytoplasm undergoes active streaming coupled to cell motility, but this streaming is laminar flow which should not be effective for mixing. In this work we used a combination of live cell tracking of injected beads and computational analysis of motion and mixing in giant amoeba *Chaos carolinensis* with the initial goal of testing the possibility that large-scale cellular deformations during pseudopod formation might implement chaotic mixing by a Baker-transform like process. Instead, we found that *Chaos carolinensis* accelerates cytoplasmic mixing using a novel cytoplasmic gel state capture and release strategy. While it was previously thought that the amoeba sol to gel state transitions only occur at the trailing and leading edge of the cell body, our work indicates that these transitions occur frequently throughout the mid-cell region, driving the cytoplasmic mixing of beads and organelles. These results indicate that amoeba achieves nearly complete mixing between 1 and 2 cytoplasmic stream/flow cycle, effectively approximating the Bernoulli mixing regime and thus representing one of the theoretically fastest possible mixers.

## INTRODUCTION

The cytoplasm is a densely packed environment, where diffusion alone is often insufficient for distributing macromolecules and organelles. This limitation arises due to the crowded nature of the cytoplasm, filled with proteins, organelles, and various molecular structures that significantly hinder the movement of particles. In cells, the large size of macromolecules and organelles combined with the high viscosity of cytoplasm means that diffusion is ineffective for long-distance transport ^1^. Even the smallest organelles, like peroxisomes which range between 0.1 and 1um ^10^, have difficulty moving short distances within a reasonable amount of time. For example, in the hypha of the fungus Ustilago *maydis,* it is calculated that peroxisomes, measured to have a diameter of ∼0.23 µm, would take roughly 42 minutes to travel 25 µm through the cytoplasm using diffusion alone ^11^. Similarly in BS-C-1 mammalian cells it is reported that an increase in lysosome size from 0.52 µm to 1.3 µm results in a decrease in lysosome diffusion coefficients ^12^, reflecting the effects of the size on diffusion at such high viscosity environments. In many cases, large biological structures are formed in a location that is distant from the point at which they are required to function, and diffusion limits the ability of such structures to reach the point at which they are needed.

To overcome this limitation, some cells have evolved mechanisms to locally synthesize or secrete molecules using strategically localized organelles. This is most evident in the development of reticulated organelle networks, such as the endoplasmic reticulum (ER) and mitochondrial network, which are coordinated by association with the cytoskeleton. Diffusion limitation can also be avoided by transporting intracellular material using motor proteins which bind cargo and walk along the cytoskeleton, anchored motor proteins which generate cytoplasmic flow that can carry material from one point to another, or by directly tethering material onto an intracellular network then using its growth and regulation for distribution.

In many cell types, intracellular materials are typically transported through cytoplasmic flows, either directly through motors pulling the cargo on cytoskeletal tracks, or passively via advection of cargo within flowing cytoplasm. In particular, larger cells have developed active mechanisms to facilitate transport through bulk cytoplasmic streaming. In these cells cytoplasmic streams are generated primarily through two mechanisms: stationary cortical motor proteins that drive cytoplasmic flow or through cellular deformation that generates cytoplasmic flow through movement of the cell boundary ^2, 3^.

Large, stationary cells, such as the syncytial *Drosophila* embryo before gastrulation and plant cells, rely on stationary motor proteins to drive cytoplasmic streaming ^2^. In contrast, motile cells like neutrophils and giant amoebas create cytoplasmic streams through active membrane deformations that accompany cellular motility ^3^. Such cells presumably rely on advective transport because they lack reticulated organelle networks ^4^ to facilitate localized delivery via targeted secretion and synthesis, nor do they maintain cytoskeletal tracks of sufficient length to transport cargo across the cell. In such cells, it is apparent that advection dominates over diffusion as a transport mechanism. This can be quantified by the Péclet number (Pe) which describes the balance between advection and diffusion for an object moving at velocity *U* over a length *L*, with *D_t_*_ℎ*ermal*_ as the diffusion coefficient. The *Pe_t_*_ℎ*ermal*_ is given by:

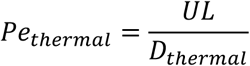

Where:

- *U* is the characteristic velocity of the fluid,
- *L* is the characteristic length over which transport occurs,
- *D_thermal_* is the diffusion coefficient expected from Brownian motion driven by thermal energy alone. It was calculated using the Stokes-Einstein equation.

In the *Chaos* genus of amoebas, nuclear size ranges from 19 µm in *Chaos carolinensis* ^13^ (used in this study), to 25.7 µm in *Chaos neos* ^14^, to 34.3 µm in *Chaos diffluens* ^14^, and 55.8 µm in *Chaos nitida* ^14^. To transport such large organelles across seemingly vast distances, amoeba *Chaos* uses membrane deformations to create advection and enhance diffusion, allowing for long distance transport of material within a reasonable time scale. Using the Stokes-Einstein equation to estimate the Brownian diffusion expected from thermal energy alone, we calculated the thermal Peclet number for 20 µm diameter beads in the sol and gel states of the cytoplasm. This analysis yielded thermal Peclet numbers of approximately 3.05 x 10^6 for the sol and 5.45 x 10^7 for the gel layers, indicating that Brownian diffusion is negligible relative to advective transport in both cytoplasmic states ^15,16,17^.

But flow is only part of the challenge. In addition to moving organelles from their site of formation to their site of use, it is also important to mix them relative to each other, in order to allow interactions and exchange of material. Given that giant amoeboid cells like *Chaos* have strong advective flows for transport, do these same flows also provide a way to mix the cytoplasm? Whether or not fluid flow causes mixing depends on the Reynolds number (Re), a dimensionless parameter in fluid mechanics that helps predict whether a fluid flow will be laminar or turbulent. It represents the ratio of inertial forces to viscous forces, providing insight into how smoothly or chaotically fluid will flow under specific conditions ^5, 6, 7, 8^. Re depends on the fluid density, flow velocity, viscosity, and the length scale of the system. For high viscosity flow in structures the size of cells, Re is extremely small. For instance, the Re for a 10 µm C. elegans pronucleus moving at 0.1 µm/s through the cytoplasm is reported to be as low as (10^-9^).^9^ Such a low Re indicates that the flow is highly laminar, with very little turbulence to promote mixing. Similarly, we calculate that the ∼20 µm diameter nuclei of the amoeba *Chaos carolinensis*, traveling at 10 µm/s, in a viscosity of 0.0045 Pa·s ^57^ (viscosity of cytoplasmic stream in *Amoeba proteus*) has a Re of approximately 4.4 x 10^⁻5^, well within the laminar flow regime.

Flow can only cause mixing at high Re, where turbulence causes different regions of the fluid to become re-arranged in an unpredictable pattern. When Re is small, fluid flows in a laminar fashion without turbulence, which means that even though the fluid is moving, neighboring parts of the fluid are all moving together, such that the relative positions of anything carried by the fluid remain in the same relative positions. We are thus left with a fundamental unanswered question – how can cells mix their contents when cytoplasmic flow is laminar?

The challenge of mixing at low Re has been investigated in several non-biological contexts. One well known theoretical model for mixing is the Baker Transform, in which a region of space containing a set of points is squashed via a continuous deformation along one axis, then split in half and the two halves stacked on each other, mimicking the way a baker folds dough ^59^ It can be shown that repeating this process leads to complete re-arrangement of the initial positions of the points, i.e. mixing, with points becoming separated as an exponential function of the number of cycles of squashing and stacking. Another type of solution is periodic alteration in the flow pattern, sometimes called “blinking flow” which allows a subset of particles to be swept in a different direction than the other particles ^58^. Again, as cycles of these alternating flows proceed, mixing gradually occurs, even though the motion is completely laminar within each flow period. In both types of models, the position of a tracer particle depends sensitively on its initial position such that its ultimate position is effectively unpredictable, thus representing “chaotic mixing” which does not require turbulent flow in the traditional sense. Given that cells can rearrange their component parts, for example by growing, moving and retracting pseudopods, and can alter the direction of their streaming flows over time, one possible solution to the mixing challenge could thus be some form of chaotic mixing. In such a model, we would expect mixing to be driven by successive rounds of pseudopod formation/retraction, and that complete mixing would be approached asymptotically over some number of cycles of cytoplasmic flow each consisting of cytoplasm flowing to the front and then returning to the rear of the cell, with the number of cycles depending on the exact nature of the chaotic mixing.

However, effective intracellular mixing likely requires additional complex flows, as seen in active turbulence, which enhances dispersion ^9^. Chaotic mixing schemes such as the baker transform rely on external forces to reshape the flow of a passive material. However, it is essential to recognize that cytoplasm is not a simple passive fluid, but an active material that can undergo state transitions. The first observations of protoplasm, consisting of the gel ectoplasm and sol (referring to solution phase) endoplasm (cytoplasmic stream), were made by Ecker in 1848¹⁸. However, it wasn’t until 1917 that Hyman proposed the sol-gel theory¹⁹. The sol-gel theory explains how amoeba controls their movement and shape changes using the transformation between two states of their cytoplasm. This idea was further advanced by Mast in a series of works in 1926, 1931, and 1934 ²⁰, ²¹, ²². Since then, modern studies have continued to probe the sol to gel transition²³, ²⁴. Most studies of the sol to gel transition have concentrated on the leading edge of the cell, where the sol state of the cytoplasm transitions into a gel state as the pseudopod extends, and at the rear of the cell, where the gel state reverts to a sol state to replenish the cytoplasmic supply at the front²⁵, ²⁶. However, because the sol to gel transition zone is not directly visible using light microscopy, except at the front and rear of the cell, its spatiotemporal dynamics and potential variation along the cell body remain poorly characterized.

A comprehensive understanding of the molecular organization and regulation of the sol to gel transition remains incomplete. It is currently hypothesized that the sol and gel layers are composed of the same cytoplasmic material, although detailed proteomic profiles of these layers are still lacking. Consistent with this idea, in vitro studies have shown that cytoplasmic extracts can transition between gel and liquid states, even demonstrating fluid streaming between the two phases in extracts ^27^. Furthermore, reconstituted systems containing actomyosin components have also replicated these properties, suggesting that it is the organizational arrangement of the cytoplasmic components, rather than their composition, that differentiates the sol and gel states^28^.

Historically, light microscopy revealed regions of actin enrichment in amoeba; however, the technology at the time was limited in its ability to visualize the organization of actin filaments at the microscale ^29^. To observe the microscale organization of filaments many studies turned to transmission electron microscopy (TEM) which enabled detailed visualization of cellular regions known to exist in either sol or gel states. For instance, studies comparing the hyaline cap of the leading edge—characterized by sol state cytoplasm—with the surrounding gel layer revealed striking differences: the sol state cytoplasm contains sparse, thin actin filaments, while the gel layer exhibits both thin and thick filaments arranged into a dense actin network ^30, 31^. Intriguingly, these filament structures have also been reconstituted in vitro from cytoplasmic extracts under conditions that induce either relaxation or contraction, thereby mimicking the native sol to gel transitions observed within the cell ^32, 33, 34^. While studies employing electron microscopy based imaging have revealed organizational differences between actin filaments in sol and gel states, they lack the ability to capture transitions between sol and gel states due to their need for fixed samples. One current question is where within the cell such transitions occur.

Here we show the amoeba *Chaos carolinensis* uses the sol-gel transition to achieve near complete cytoplasmic mixing within 1-2 flow cycles. where we define a flow cycle as the time taken for a particle to traverse the full length of the cytoplasmic sol layer and return via the cytoplasmic gel layer. This approximates the Bernoulli mixing regime, in which the position of all particles in a region of material is completely randomized in one step ^60^. This mixing thus occurs much more rapidly than could be explained by chaotic mixing models which, as noted above, typically approach full mixing asymptotically over many cycles.

Our findings indicate that *Chaos carolinensis* achieves such rapid mixing by inducing sol to gel transitions within the mid-cell region, effectively transferring particles, such as beads and nuclei, between the sol and gel states. This dynamic enables particle exchange across the cytoplasm layers, promoting mixing. We explore this mechanism here by simulating cytoplasmic flow including and excluding mid-cell state switching and comparing mixing behaviors. We validate our simulation by examining the mixing of two bead populations at different time delay intervals, using a method we developed to computationally label bead populations based on tracking data from a single experimental label. Analysis of particle velocities as they approach the sol-gel boundary suggests that the mixing is not simply an effect of a no-slip boundary condition, but involves discrete capture events by the gel. These insights not only broaden our understanding of amoeboid cytoplasmic dynamics but also suggest that sol-gel transitions may represent a generalizable mechanism for efficient intracellular mixing in cells with cytoplasmic streaming.

## RESULTS

### Experimental Mixing of Microinjected Beads Across Delays

To observe intracellular mixing, we microinjected fluorescent beads into the amoeba *Chaos carolinensis* and performed time-lapse microscopy combined with a custom image-analysis pipeline. This pipeline splits the cell into two halves and assigns beads to two groups based on which half of the cell they were in, then tracks them, allowing their motion to be analyzed as two distinct, virtually labeled bead populations from a single injected fluorescent species. This approach enables the analysis of mixing dynamics starting from any frame within a time-lapse acquisition, using just a single population of injected beads. Using this approach, we quantified mixing across variable temporal delays with a Simple Nearest Neighbor (SNN) metric, which measures the spatial intermixing of two bead populations over time by counting the number of closest neighbors of the same versus different colors to produce a score that varies from 0 for completely separated colors to 0.5 for completely mixed colors (see Methods, Formula 2).

Traditionally, SNN-based analyses require microinjection of two spectrally distinct bead populations, a technically demanding procedure that necessitates simultaneous on-scope injections and restricts measurements to a single mixing event following injection **(Fig. 1.1)**. To overcome these constraints, we developed an alternative strategy in which a single bead population is digitally partitioned into two artificially colored groups during post-processing. This method permits initiation of mixing analyses at any point in the time series, if bead trajectories remain continuous, thereby enabling repeated and temporally flexible measurements within the same dataset **(Fig. 1.2 and Fig 1.3)**.

**Figure 1.**
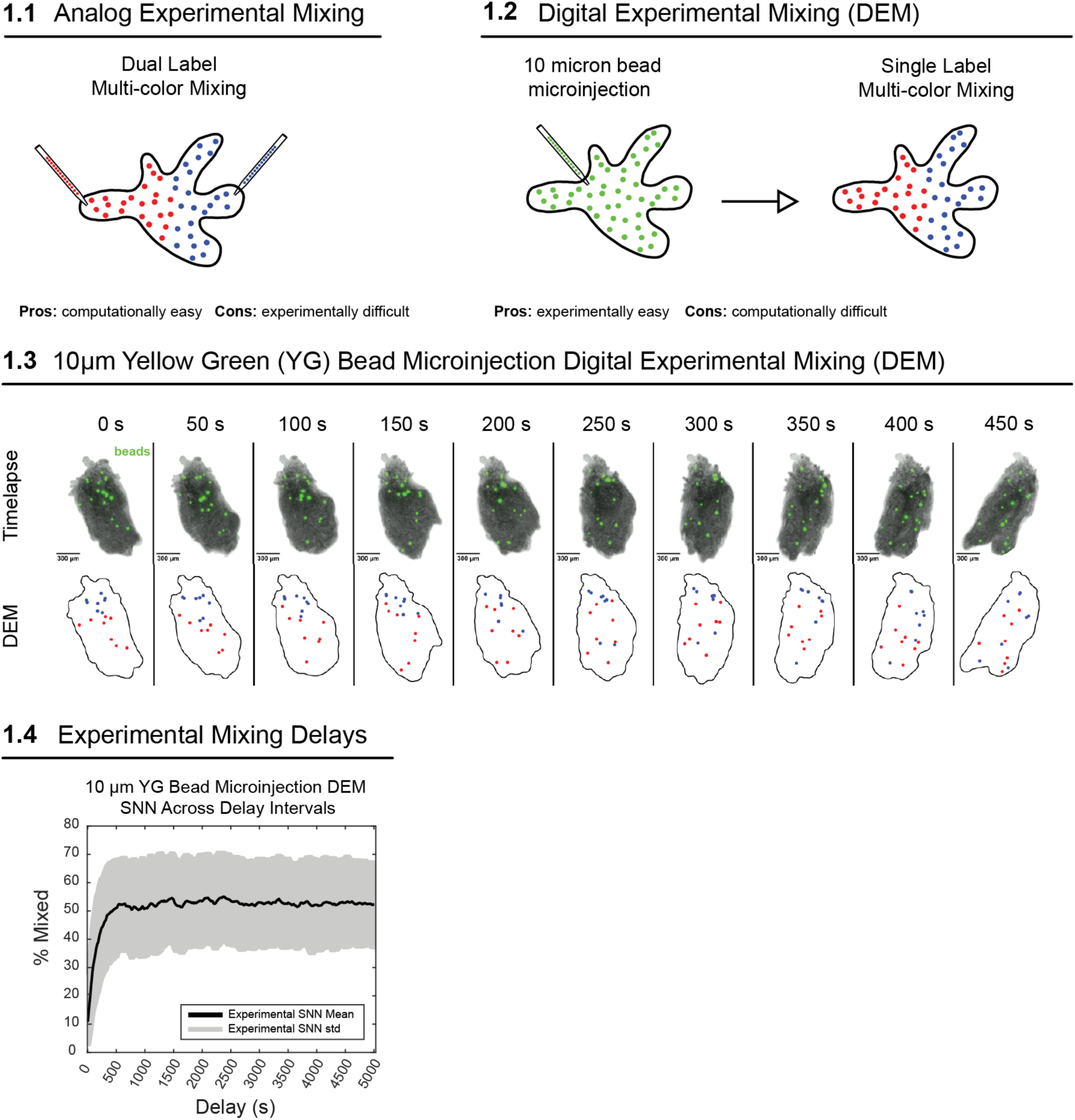
Experimental Mixing of Microinjected Beads Across Delays. 1.1 Analog experimental mixing measurement by injecting beads of different colors. This method is computationally easy but experimentally difficult. 1.2 Digital experimental mixing (DEM) using 10 µm bead microinjection. This method uses a single label combined with virtual labelling and tracking to establish multi-color mixing. This method is experimentally easy but computationally challenging. 1.3 Time-lapse images showing the digital experimental mixing (DEM) process using 10 µm YG bead microinjection. Bead movement is tracked over 450 seconds, and the bottom row illustrates single label multi-color mixing in corresponding frames 1.4 DEM SNN calculations across time delays. The plot shows the percentage mixed as a function of delay, with mean and standard deviation indicated.

Tracking beads continuously over long periods, up to 12 hours in this case, becomes increasingly difficult when using large populations. For this reason, we only microinjected 18, 10 µm YG beads into our sample. Despite our reasonable bead population size, we generated 54 total tracks, indicating that some tracks failed to maintain continuity and started new tracks to preserve accuracy. To overcome this limitation, we developed a custom MATLAB script to filter through the 54 tracks to find video segments containing 18 continuous tracks. We then analyzed those individual video segments.

The longest continuous segment, consisting of 15,000 frames, captured 2 hour and 5 minutes of usable intracellular dynamics. From this segment, we calculated SNN scores across 10,000 mixing delays ranging from 0 to 5,000 seconds and plotted the results **(Fig. 1.4).** Complete mixing, as judged by the SNN metric reaching its asymptotic value, was achieved within approximately 440 seconds in this cell. In a later section we estimate a simulated cycle time of 220 seconds (**Fig. 6.4**).

Importantly, this time scale is far faster than would be expected from diffusion alone. For particles of this size (10 µm diameter), effective diffusion coefficients in cytoplasm are extremely small due to molecular crowding, viscoelasticity, and interactions with cytoskeletal networks. Even under optimistic assumptions, purely diffusive transport over cellular length scales on the order of 100–200 µm would require many hours to days. The rapid approach to complete mixing observed here therefore cannot be explained by Brownian motion and instead requires active, flow-mediated transport.

Given that laminar flow during cytoplasmic streaming cannot cause mixing, and given that mixing by diffusion would be far too slow, what process can account for the short mixing timescale that we observe?

### Close Bead Pairs Separation Events

While mixing is often thought of in terms of scrambling a whole continuous region of space, its effects can be observed in terms of pairs of particles embedded in that space. During mixing, particles that start out close together become far apart and vice versa, and the faster this happens, the faster the mixing. The mathematical form of the dependence of distance on time can provide a way to classify intracellular mixing with respect to known classes of mixing processes ^60^, which could then provide potential insight into the type of mechanism that might be at work. To take this type of approach, we quantified the time-dependent separation of initially neighboring tracer beads under cytoplasmic flow. To accomplish this, we microinjected two fluorescent 20 µm beads into cells and imaged their motion using time-lapse microscopy **(Fig 2.1).** To best isolate bead motion from overall cellular movement, we developed an image-processing pipeline (OttoPipeline) that transforms time-lapse videos of freely crawling cells into a stationary crawling reference frame, as described in the Methods **(Supplementary Fig. 4)**.

**Figure 2.**
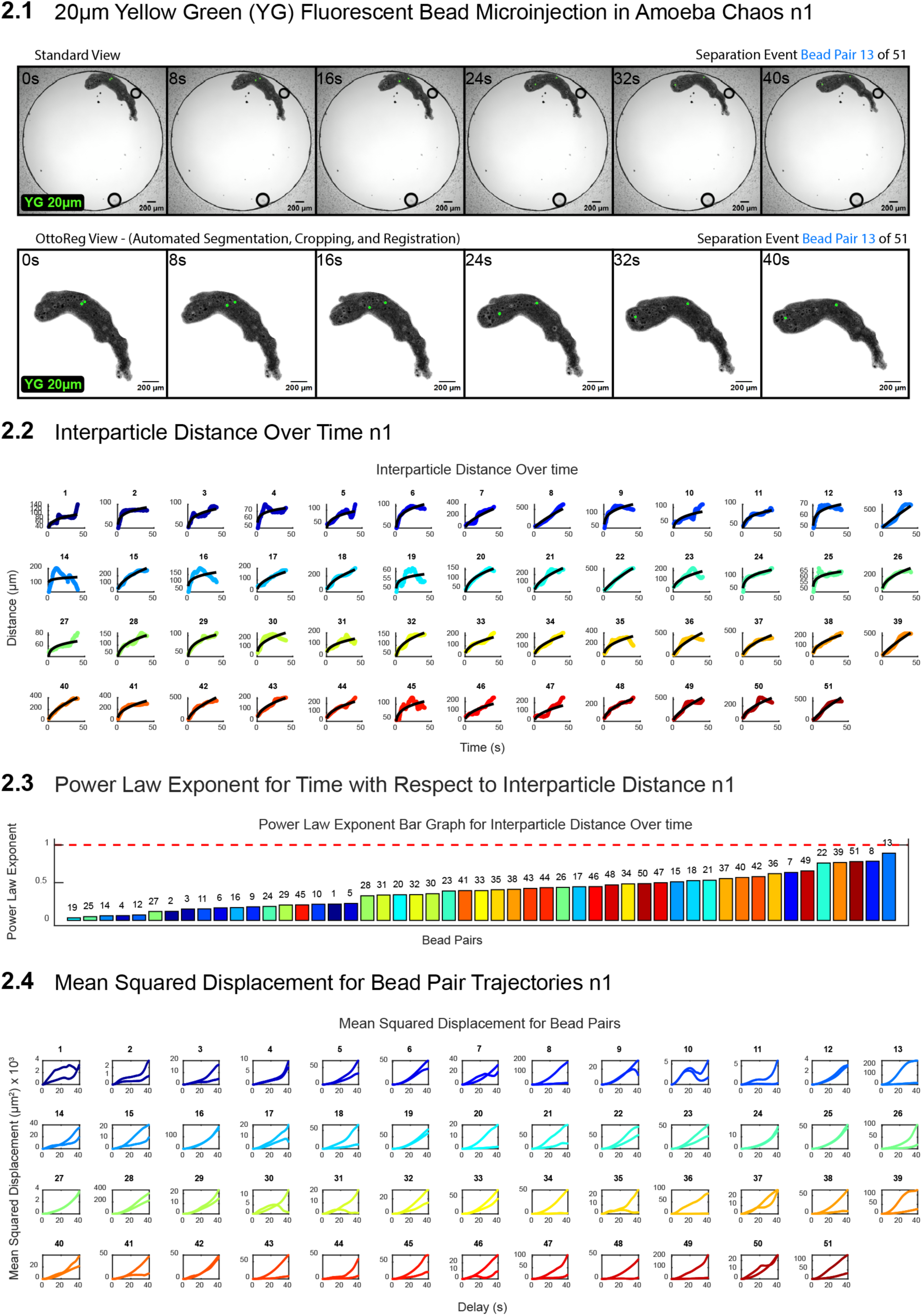
20 µm Bead Separation Assay n1 of 4. 2.1 Time-lapse images showing the microinjection of 20 µm yellow-green (YG) fluorescent beads into amoeba *Chaos*. The top row presents the standard view, and the bottom row shows the OttoReg view, where automated segmentation, cropping, and registration have been applied. The images capture a separation event for bead pair 13. Scale bars represent 200 µm. 2.2 Interparticle distance over time for each bead pair tracked throughout the experiment. Each plot corresponds to a bead pair, displaying the distance (µm) between beads over time (s). 2.3 Power law exponent for interparticle distance as a function of time. The bar graph shows the power law exponent for each bead pair, with values color-coded and labeled. The red dashed line is placed at Y = 1. 2.4 (MSD) plots for each bead pair trajectory. These plots show the MSD (µm²) as a function of delay time (seconds), providing insight into the diffusion behavior of the beads within the amoeba.

Mixing efficiency can be quantified by fitting a power-law exponent to the separation of initially neighboring particles over time. For this purpose, we calculate the geodesic distance between particles, i.e. the shortest path between the particles that is contained inside the cell. We use this form of distance to handle cases where two particles end up in different pseudopods such that the standard through-space distance does not reflect their separation within the cytoplasm. We then ask how this geodesic distance changes over time. In many mixing processes, particle separations grow as a power law in time, with higher exponents indicating faster and more efficient mixing, while lower exponents reflect restricted transport. This type of scaling analysis has been used extensively in studies of turbulent dispersion and, more recently, in cellular biophysics to distinguish between distinct mixing regimes ^46, 53, 54^.

In idealized laminar flows, such as those observed during cytoplasmic streaming, particles move in smooth, ordered layers. In this context, our power-law analysis describes bead-pair separation over time, and therefore measures pair dispersion, also referred to as relative dispersion, rather than the displacement of a single particle from its starting position. Laminar systems typically exhibit lower mixing efficiency than turbulent or chaotic flows because particle motion is steady and predictable, leading to regular, non-chaotic relative dispersion. Although laminar flows can transiently display linear growth in interparticle distance, resembling ballistic pair separation, they generally lack the repeated stretching and folding required for efficient mixing. For example, in boundary layers along flat plates under laminar flow conditions, nearby fluid particles can experience approximately constant shear and separate at an approximately constant relative velocity over short timescales ^55^. In this regime, bead-pair separation can increase nearly linearly with time before deviating due to diffusion, spatially varying flow, or instabilities, corresponding to a power-law exponent near 1. This behavior is characteristic of ballistic-like relative dispersion, where the distance between two particles grows linearly because their relative velocity is approximately constant ^41,42,48,49^.

Outside of this near-boundary or constant-shear regime, laminar flows are more commonly associated with slower relative dispersion. In such cases, bead-pair separation may fall within the sub-diffusive-like regime, with exponents between 0 and 0.5 ^37,43,44^, reflecting slow and inefficient separation between particles ^45^. For a purely diffusive relative-dispersion process, the pair-separation distance is expected to scale with a power-law exponent of approximately 0.5^37,38,43,44^.

Exponents between 0.5 and 1 indicate super-diffusive-like relative dispersion, meaning that bead pairs separate faster than expected from purely diffusive relative motion but slower than ballistic separation. This regime can occur in systems with active, heterogeneous, or fluctuating flows, including active turbulence at low Reynolds number ^9^. Importantly, this classification refers to the scaling behavior of pair separation and does not imply inertial turbulence. In contrast, super-ballistic relative dispersion, defined by power-law exponents greater than 1, indicates faster-than-linear growth in interparticle distance. Such behavior can occur when the relative velocity between particles increases over time, as in shock-wave-driven or explosive mixing systems ^42,50^, or in classical turbulent pair dispersion, where Richardson-type scaling predicts accelerated pair separation ^46,47^. These systems can exhibit high mixing efficiency because particle pairs separate rapidly across space. By contrast, slower mixing systems may show restricted relative dispersion except in specific regions where local geometry or flow structure promotes rapid separation. For example, the Baker’s transformation can produce super-ballistic-like relative dispersion in the localized region where cutting occurs post stretching and prior to folding.

In our data, power-law fits to the geodesic interparticle distance revealed a broad distribution of bead-pair separation behaviors. We measured the interparticle distance for 169 bead-pair separation events from four experimental samples at 25 °C **(Fig 2.2; Supplementary Figs. 1.2, 2.2, 3.2)** and fit a power-law exponent describing interparticle distance as a function of time for each event, observing similar distributions across all samples **(Fig. 2.3; Supplementary Figs. 1.3, 2.3, 3.3)**. Of these events, 7% (12 events) were super-ballistic, exhibiting a power-law exponent slightly greater than 1; 41% (69 events) were super-diffusive, with exponents between 0.5 and 1; and 50% (84 events) were sub-diffusive, with exponents between 0 and 0.5. An additional 2% (4 events) exhibited slightly negative exponents, reflecting bead pairs that moved closer together by the end of the assay due to trajectories colliding with the cell boundary.

These power-law exponents classify pair-level mixing regimes, capturing how rapidly two nearby beads separate over time as a consequence of the underlying cytoplasmic dynamics. Most bead-pair separation events fell within the super-diffusive regime, with only a small fraction entering the super-ballistic regime. The presence of super-diffusive movement was unexpected, as we did not observe turbulent flows or explosive cellular events that would readily explain such rapid pairwise separation.

To resolve this apparent discrepancy, we next examined the single-particle transport behavior underlying these pair-level mixing regimes by performing a rolling mean-squared displacement (MSD) analysis on individual bead trajectories within separating bead pairs **(Fig 2.4).** This analysis allowed us to classify time-resolved segments of individual trajectories as sub-diffusive, diffusive, or super-diffusive. Focusing on bead pairs classified as super-diffusive or super-ballistic at the pair level, we found that 63% (51/81) of these events consisted of one bead exhibiting super-diffusive motion relative to the reference frame of the cell while the paired bead displayed sub-diffusive or restricted behavior. This asymmetry suggests that one bead was advected within the sol phase, while the other remained trapped in the gel phase. In contrast, 37% (30/81) of super-diffusive and super-ballistic separation events involved both beads following super-diffusive trajectories, consistent with both beads being transported within the sol phase, potentially along divergent flow paths such as separate pseudopods. Alteration of flow pattern has been shown to drive mixing in non-biological systems ^58^.

### Pseudopod Modulation and Separation Rate

The differences in bead pair motion observed in the previous section may reflect situations in which beads become separated by entering different pseudopods which then undergo different movements. We speculated that pseudopod formation, migration, and resorption might drive mixing in a manner analogous to the Baker’s Transformation, in which a region of space is stretched and folded repeatedly. The transformation is so named because its action resembles the folding and stretching of dough by a baker. It is widely used in dynamical systems theory to illustrate how iterative stretching and folding operations lead to efficient mixing in confined domains, and it remains a classic example in the study of chaotic dynamics ^48, 52^. In theory, complete mixing requires infinitely many iterations, but in practice near-complete mixing is often achieved within 5–10 cycles ^48, 52^. We hypothesized that separation of cytoplasm into distinct pseudopods followed by pseudopod migration and resorption might produce a mixing process like the Baker transform in which the two pseudopods would correspond to the two pieces of dough. If that was the case, then entry of the two beads into separate pseudopods followed by divergence in pseudopod trajectories would produce faster separation between the beads than shared pseudopod entry by both beads. Thus, we reasoned that an increase in pseudopod number should elevate the frequency of ballistic or super-diffusive separation events, while a reduction in pseudopod number should correspondingly reduce bead separation.

To test this idea, we used non-chemical perturbations to modulate pseudopod number in microinjected bead pair time-lapse experiments. We compared the power law exponents across three experimental groups: water plus methyl cellulose at 25 °C, to increase pseudopod number; water at 37 °C, to decrease pseudopod number; and water at 25 °C as the control (*Chaos* is a pond-dwelling organism that lives in fresh water), reused from the previous section (**Fig. 3.1**). To assess the effectiveness of these perturbations, we monitored convex cell curvature across every time-lapse acquisition (**Fig. 3.2**). Our analysis showed a significant difference in the convex sum, defined as the sum of convex node angles along the cell boundary across all frames (**Fig. 3.3**), confirming the effectiveness of our pseudopod-modulating conditions. Despite observing a significant increase in pseudopod number in the methyl cellulose at 25 °C group compared to the water at 25 °C control, we found no significant difference in the power law exponents of bead separation between these groups which produced mean exponents of 0.45 (MC 25 °C) and 0.50 (water 25 °C) (**Fig. 3.5**), suggesting that increasing pseudopod number has minimal effect on ballistic or super-diffusive separation events and by extension the overall mixing outcome. In contrast, we observed a slight but significant decrease in exponents between the water at 25 °C control, with a mean exponent of 0.50, and water at 37 °C group, with a mean exponent of 0.36. This finding is consistent with the hypothesis that separation is accelerated, in part, by beads entering different pseudopods, as proposed in the previous section to explain rare super-diffusive events. At 37 °C, where pseudopod number is reduced, this class of events is diminished, leading to a modest decrease of the mean exponent. However, ballistic separations are not removed entirely at 37 °C, highlighting the need for alternative mixing mechanisms.

**Figure 3.**
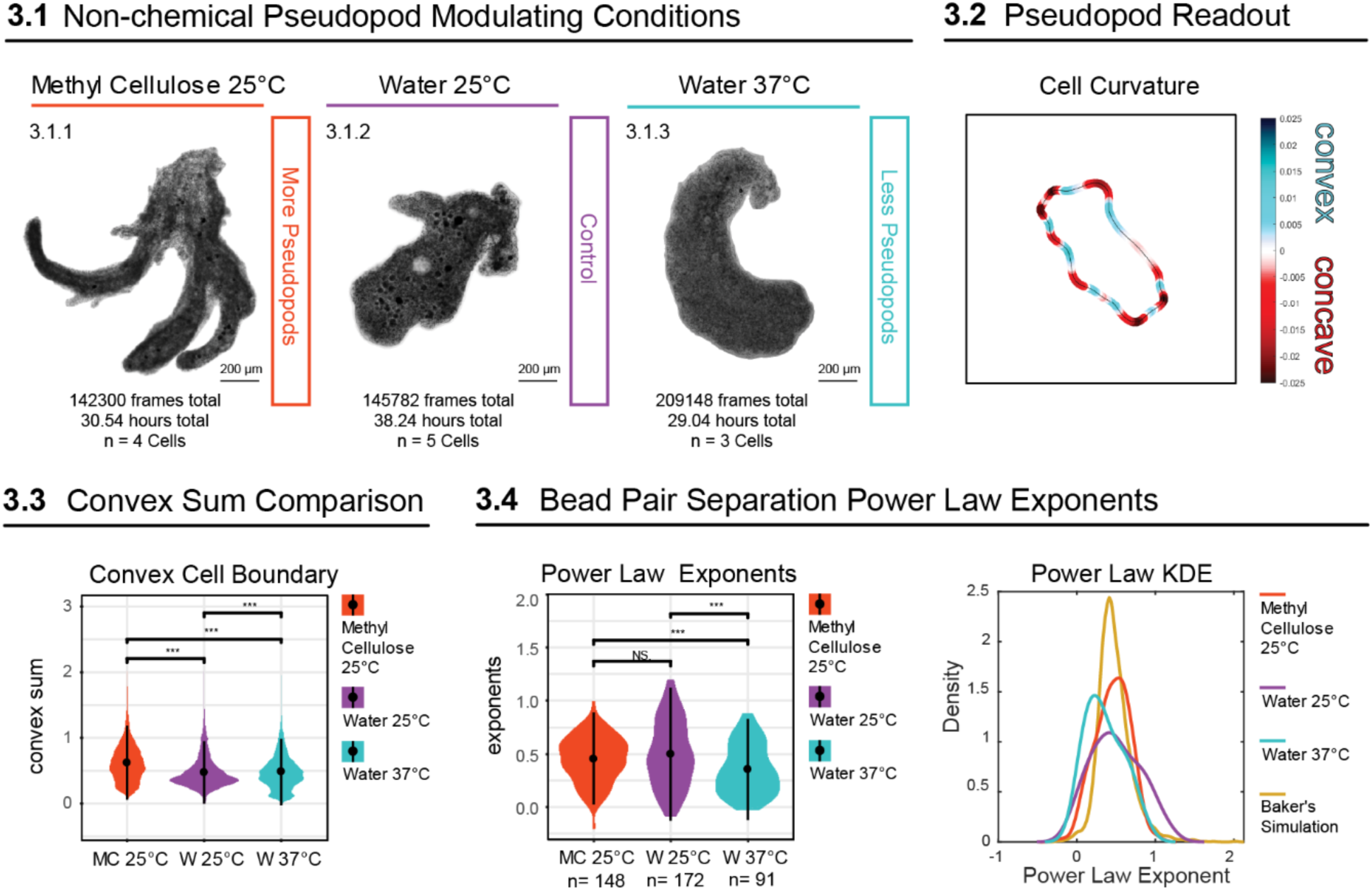
Bead Separation Rate and Pseudopod Number. 3.1 Non-chemical pseudopod modulating conditions: 3.1.1–3.1.3 Time-lapse images showing amoeba in three different environmental conditions: Methyl Cellulose at 25 °C, Water at 25 °C, and Water at 37 °C. The methyl cellulose condition at 25 °C increases pseudopod number whereas the water condition at 37 °C reduces pseudopod number. The water condition at 25 °C represents normal number of pseudopods. The total number of frames, hours recorded, and number of cells (n) are indicated. 3.2 Pseudopod number readout: Cell curvature analysis is shown, where convex (blue) and concave (red) regions along the cell outline correspond to areas where pseudopods are found. 3.3 Convex sum comparison: Violin plot showing the distribution of convex sums under different environmental conditions (Methyl Cellulose 25 °C, Water 25 °C, and Water 37 °C). The asterisks indicate statistically significant differences between conditions, marking the effectiveness of pseudopod modulation under these conditions. 3.4 Bead pair separation events: Scatter plots showing the position of bead pairs under the three conditions at time 0 seconds (top row) and 40 seconds (bottom row). The number of bead pairs (n) for each condition is indicated. 3.5 Power law exponents: Violin plot showing the separation rate (power law exponent) for bead pairs across the three environmental conditions. Significant differences between conditions are indicated by asterisks, with “NS” indicating non-significant differences.

Power law exponents describing nearest-neighbor separation were calculated for both experimental data and a Baker’s Transformation simulation to provide direct comparisons between the two. In the experimental dataset, the distribution of exponents was centered near α≈0.5. Applying the same analysis framework to the Baker’s Transformation model (fixed nearest neighbors at t=0, cycle-boundary sampling, and whole-trajectory log–log fits) produced a similar unimodal distribution with mean α=0.48 (median = 0.45). As shown in **Figure 3.5**, the experimental and simulated bakers transformation distributions overlap closely, supporting the conclusion that the Baker’s Transformation may capture some key features of particle separation dynamics in living cells. However, while the Baker’s Transformation is a powerful tool for chaotic mixing and enhances the speed of homogenization, it is not necessarily associated with super ballistic mixing observed in the control water at 25 °C. The presence of super ballistic transport depends on the specific features of the flow and the existence of regions or mechanisms that facilitate this anomalous diffusion ^56^, such as at the border where cutting/stacking/folding occurs after stretching in the bakers transform. Our control results show an increase in these kinds of ballistics events compared to the bakers transform, suggesting that there may be many multiple regions of anomalous diffusion within the amoeba *Chaos* system.

### PIV Observations of Sol and Gel Layers

The fact that cases of ballistic separation generally entailed one bead moving while another remains approximately stationary raised the possibility that some mechanism might transiently arrest the motion of some sub-set of beads. The morphology of the amoeba consists of two layers of gel sandwiching a layer of sol state cytoplasm that streams via a mechanism involving the selective blebbing of the actin cortex at the leading edge. The leading edge maintains a continuous sol to gel state transition. Meanwhile, the rear of the cell, known as the uropod, maintains a continuous gel to sol state transition ^18, 19, 20, 21, 22^ **(Fig 4.1)**. Within this sandwich structure, the gel phase is fixed with respect to the lab reference frame, so that any object within, or adhering to, the gel phase would show little or no motion. This raises the possibility that super-diffusive bead motion might involve transient capture of one bead by the gel cortex, with the other bead moving in the sol phase flow.

**Figure 4.**
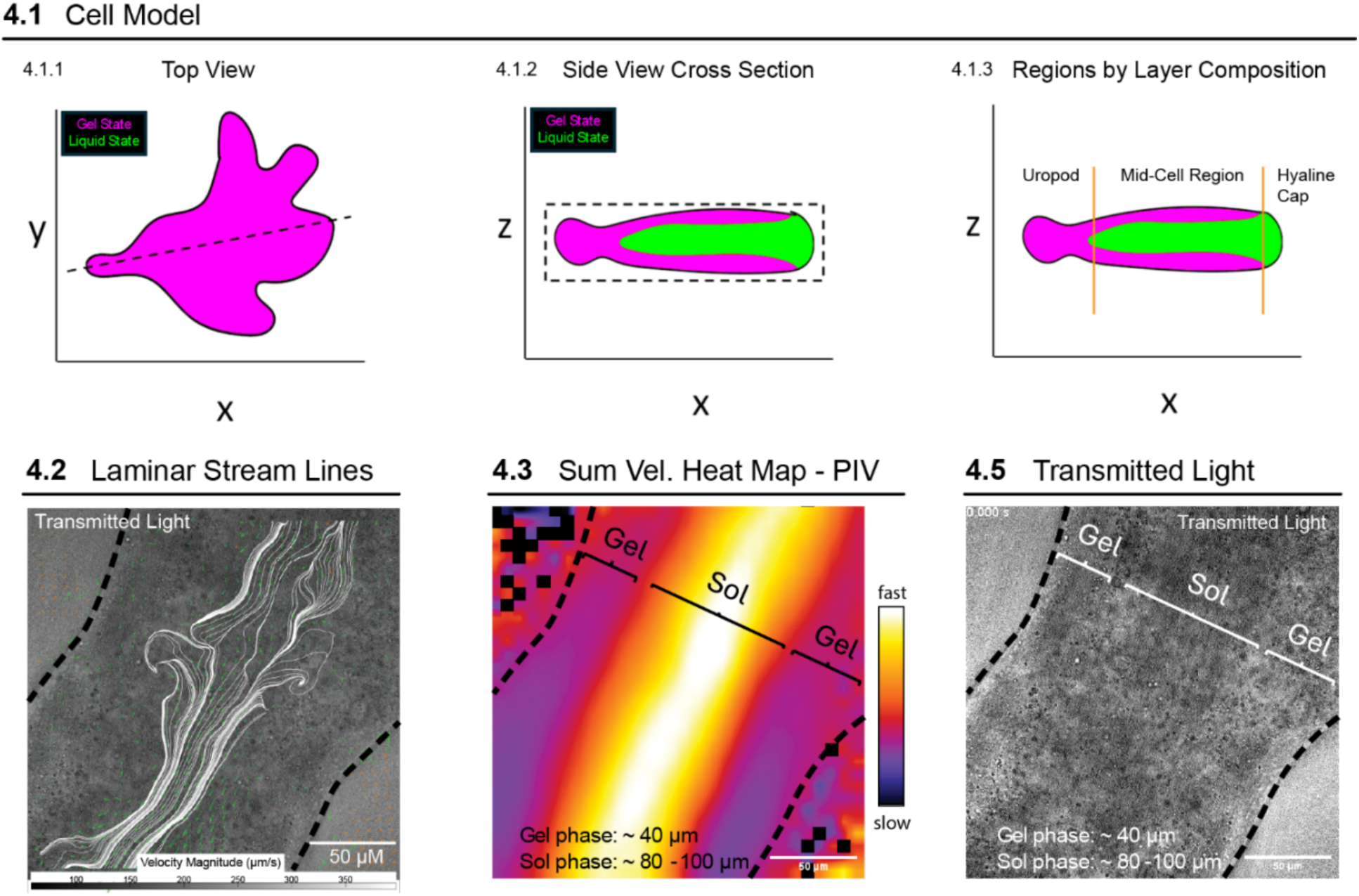
PIV Observation of Gel and Sol State Flow Layers. This figure illustrates the morphological distribution of gel and liquid states within an amoeba. 4.1 Cross-sectional view in the x-y plane highlights the overall shape of the amoeba. 4.2 Side view (x-z-plane) shows the stratification of states along the length of the cell. The dashed line marks the axis of view for the side projection, with the liquid state concentrated in the central core, surrounded by the gel state. The dashed line marks the axis of view for the side projection. This highlights the spatial dynamics of cytoplasmic sol to gel transitions in amoeboid movement. 4.3 Laminar flow streamlines obtained from particle image velocimetry (PIV) analysis in the liquid (sol) state of the cytoplasm. The streamlines illustrate the smooth, parallel flow of cytoplasmic contents in this region. The dashed line indicates cell boundary. The flow velocity is measured in microns per second (µm/s), with a scale bar representing 50 µm. 4.4 Heat map showing the sum of velocity magnitudes in the sol and gel phases of the cytoplasm. The warmer colors represent faster velocities, while cooler colors correspond to slower velocities. Gel and sol regions are marked, showing a clear difference in flow characteristics across these mid-cell regions. 4.5 Transmitted light image showing the distinct separation between gel and sol regions in the amoeba’s cytoplasm. The boundaries are labeled accordingly, and the gel phase (∼40 µm) and sol phase (∼80–100 µm) thicknesses are noted. The dashed lines mark the cell boundary.

As a first step towards testing this idea, we first measured the geometry of the two phases. While gel and sol states are easily distinguishable at the front and the back of the cell, we sought to provide evidence for laminar flow in the sol state layer and to determine the thickness of sol and gel state layers at the mid-cell region where the difference between layers is not as visually obvious. To search for laminar flow and the border between sol and gel state layers we turned to Particle Image Velocimetry (PIV). PIV is an optical measurement technique used to measure fluid flow by tracking the movement of particles within fluid. PIV adopts an Eulerian approach, focusing on fixed points within a flow field and measuring how the fluid moves through these points over time. This contrasts with a Lagrangian approach, which would track individual particles as they move through the flow, as we have done in previous sections. By capturing velocity information across a grid of fixed points, PIV provides a comprehensive map of flow patterns. This makes PIV particularly useful for studying complex, spatially distributed flow fields. Using PIV, we traced the bulk flow of cytoplasmic vacuoles, vesicles, and refractile bodies observed from transmitted light timelapse acquisitions. We detected laminar flow streamlines in cytoplasmic streams **(Fig 4.2)**, confirming our assumption that cytoplasmic streams are laminar. We used the sum of velocities, u and v components, to find interfaces between different flows, which effectively marks the borders between sol and gel states in the mid-cell region **(Fig 4.3)**. From these measurements we note the gel phase is roughly 40 µm on either side with the sol phase being 80 – 100 µm. Without PIV analysis the border between sol and gel states would be visually indistinguishable in this region of the cell **(Fig 4.4)**.

### Gel and Sol State Flow Properties

Having confirmed that both gel and sol layers exist all along the cell, we considered the possibility that adhesion to, or entrapment in, the gel layer of particles travelling in the sol layer might cause some particles to pause, as observed in the bead motion data. To explore this idea, we mapped bead transitions into each layer (sol versus gel) along the normalized cell length using our control group (water, 25 °C) data from previous sections. We first segmented and classified flow states within individual trajectories. To do this, we applied a rolling mean squared displacement (MSD) analysis and assigned a mean MSD value to each time point **(Figs. 5.1 and 5.2)**, enabling classification of state changes throughout the time-lapse data. Flow states were then separated using a manually selected threshold, which was fine-tuned to group periods during which beads exhibited distinct modes of motion. Fine tuning was done by overlaying bead trajectories on videos with color-coded state labels, green for sol flow and magenta for gel flow, allowing visual validation of classification accuracy. This was possible because state changes are visually distinct by their apparent reversal in flow. **(Fig. 5.2)**.

**Figure 5.**
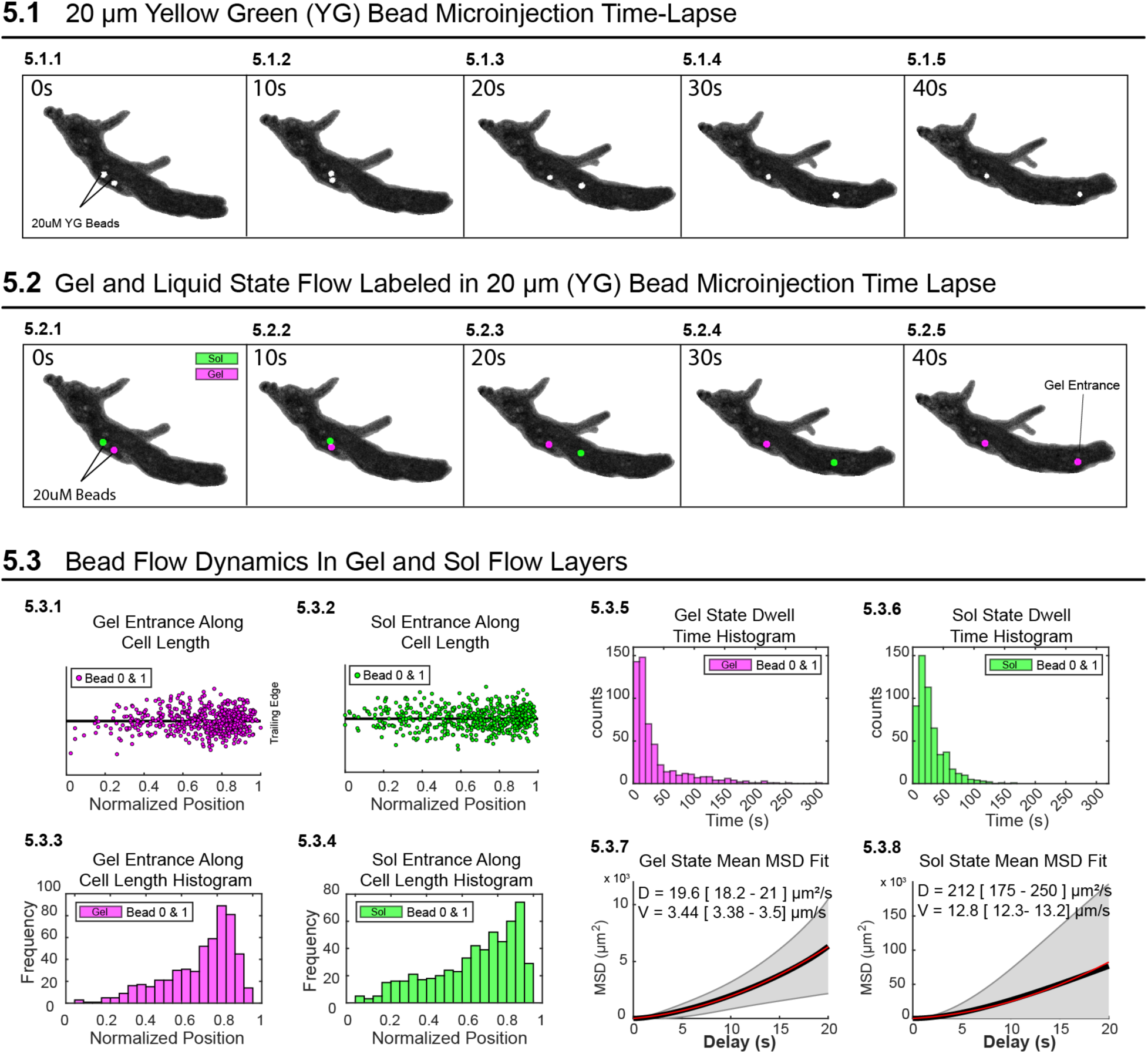
Gel and Sol State Flow Properties. 5.1 Time-lapse images of 20 µm Yellow Green (YG) fluorescent bead microinjection into an amoeba. Bead movement is tracked over time, with images captured every 0.5 seconds from 0 to 40 seconds. 10 second intervals between frames are shown in this panel. 5.2 Gel state (magenta) and Sol State (green) overlayed onto bead timelapse. 5.3.1–5.3.2 Gel state (magenta) and Sol State (green) normalized location transition sites along cell length. 5.3.3–5.3.4 Histograms of normalized location transition sites along cell length. 5.3.5–5.3.6 Histograms of dwell times for the gel and liquid states for Beads 0 and 1. 5.3.7–5.3.8 MSD fits for gel / liquid states. Effective diffusion coefficients (D) and mean velocities (V).

This approach also allowed us to analyze the properties of each layer separately to measure dwell time distributions, diffusion coefficients, and mean velocities of beads flowing in either layer using the apparent diffusion constant as an indicator of whether a bead was in the sol or gel state.

Our results indicate that both sol and gel entrance events are enriched toward the front of the cell **(Fig. 5.3.1 – 5.3.4)**, contrary to traditional models of rear-biased gel to sol state transition. From MSD curve fits, we obtained diffusion coefficients of 19.6 µm²/s for the gel state **(Fig. 5.3.7)** and 212 µm²/s for the solution state **(Fig. 5.3.8)**. To disentangle the relative contributions of diffusion and advection, we estimated the diffusion coefficient expected under purely diffusive conditions using the Stokes–Einstein equation as a theoretical baseline:

**Using the Stokes-Einstein relation**

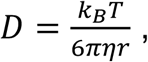

with:

- *k_B_* = 1.38 × 10^−23^ J/K
- *T* = 310 K
- *r* = 10 × 10^−6^ m (10 µm radius)
- *η* = viscosity

**Ectoplasm (moderate viscosity)**

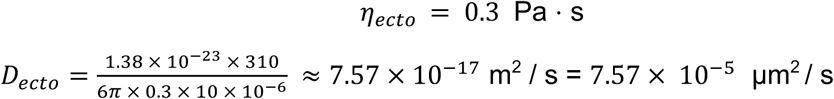

**Endoplasm (low viscosity)**

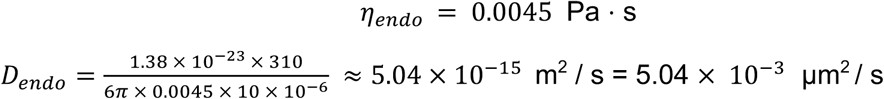

Direct viscosity measurements are not available for *Chaos carolinensis*, so we used previously reported measurements for the ectoplasm, corresponding to the gel state, and endoplasm, corresponding to the sol state, of *Amoeba proteus* ^57,61^. Using viscosities of 0.3 Pa s for the gel state ^61^ and 0.0045 Pa s for the sol state ^57^, we used the Stokes-Einstein equation to estimate the Brownian diffusion expected for 20 µm beads in each cytoplasmic state. This yielded predicted thermal diffusion coefficients of 7.57 x 10^-5^ µm^2^/s in the gel state and 5.04 x 10^-3^ µm^2^/s in the sol state. These values provide an estimate of the bead motion expected from thermal diffusion alone and indicate that passive Brownian motion should be negligible on the length scale of the cell.

However, the measured diffusion coefficients from bead dispersion in *Chaos* were several orders of magnitude higher than these Stokes-Einstein predictions. We measured effective diffusion coefficients (*D_effective_*) of approximately 20 µm^2^/s in the gel state and 212 µm^2^/s in the sol state. These measured values cannot be interpreted as thermal diffusion coefficients. Instead, they represent effective or active dispersion coefficients that include contributions from cytoplasmic streaming, flow heterogeneity, and active cytoplasmic remodeling. Therefore, to avoid conflating Brownian diffusion with experimentally observed bead dispersion, we distinguish between a thermal Peclet number and an active Peclet number.

The thermal Peclet number, calculated using the Stokes-Einstein diffusion coefficient, quantifies the relative importance of advective transport compared with Brownian diffusion alone. Using the measured bead velocities and a characteristic cell length of 1200 µm, we calculated thermal Peclet numbers of approximately 5.45 x 10^7 for the gel and 3.05 x 10^6 for the sol. These very large values indicate that Brownian diffusion is negligible relative to advective transport in both cytoplasmic states.

In contrast, the active Peclet number was calculated using the experimentally measured effective diffusion coefficients. The active Peclet number, *Pe_active_*, is given by

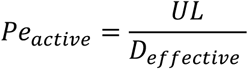

Where:

- *U* is the characteristic velocity of the fluid,
- *L* is the characteristic length over which transport occurs,
- *D_effective_* is the diffusion coefficient calculated from experimentally measured MSD curves from bead trajectories.

This value does not compare advection to thermal diffusion but instead compares directed cytoplasmic transport to the observed active dispersion of beads in the living cell. Using the measured bead velocities and effective diffusion coefficients, we calculated active Peclet numbers of approximately 210.61 for the gel state and 72.45 for the sol state. Both values are greater than 1, indicating that bead motion is advection-biased in both cytoplasmic states. However, the higher active Peclet number in the gel should not be interpreted as faster flow. Rather, it reflects that bead motion in the gel is more directionally constrained relative to its lower measured dispersion. Conversely, the lower active Peclet number in the sol reflects rapid directed flow accompanied by greater bead dispersion. Together, these two Peclet numbers show that Brownian diffusion is insufficient to explain bead motion, while active cytoplasmic processes strongly enhance particle dispersion beyond thermal expectations.

Moreover, our dwell-time analysis **(Figs. 5.3.5 and 5.3.6)** shows that most sojourns in either state last approximately 20 seconds, with many lasting under 10 seconds, implying frequent transitions between layers before beads reach the leading or trailing edges of the cell. To further assess whether diffusion alone could explain our observed transitions, we estimated the characteristic diffusion-to-capture time for a bead to traverse a sol layer using the mean first-passage time for a uniform initial distribution in a slab with both walls absorbing given by

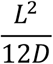

where:

- *L* is the characteristic length in the system: sol layer thickness
- *D* is the diffusion coefficient of a particle or molecule in the medium: Here we use *D_effective_* for *D*.

This approximation yields a diffusion time of 3.93 seconds for a 100 µm thick sol layer indicating that diffusion within the sol phase is fast enough to account for the observed time spent in that phase. However, this also suggests that particles encountering the sol–gel boundary are not always immediately captured and immobilized, and that additional regulatory mechanisms, potentially including mechanical gating or cytoskeletal restructuring, control bead partitioning between flow states.

Finally, we observed that bead transitions between sol and gel layers occur not only near the leading and trailing edges, as traditionally assumed, but also throughout the mid-cell region with increasing concentrations towards the leading edge of the cell for both layers **(Figs. 5.3.1–5.3.4)**. The short dwell times **(Figs. 5.3.5 and 5.3.6)** and frequent mid-cell transitions indicate that sol–gel switching is a dynamic, spatially distributed process that occurs across the entire cell, rather than being confined to leading and trailing regions.

### Mid-Cell Sol-Gel Transitions Accelerate Mixing

To study the effects of mid-cell sol–gel state switching on intracellular mixing, we developed a simulation framework guided by the experimentally measured flow properties of the gel and sol layers described in the previous section (“Gel and Sol State Flow Properties”). In our simulation, the spatial organization of gel and sol layers reflects amoeba morphology in terms of the organization into layers and the approximate size of the cell **(Figs. 4.1–4.3)**, but is simplified into three stacked planes, each measuring 200 µm × 200 µm **(Fig. 6.1)**.

**Figure 6.**
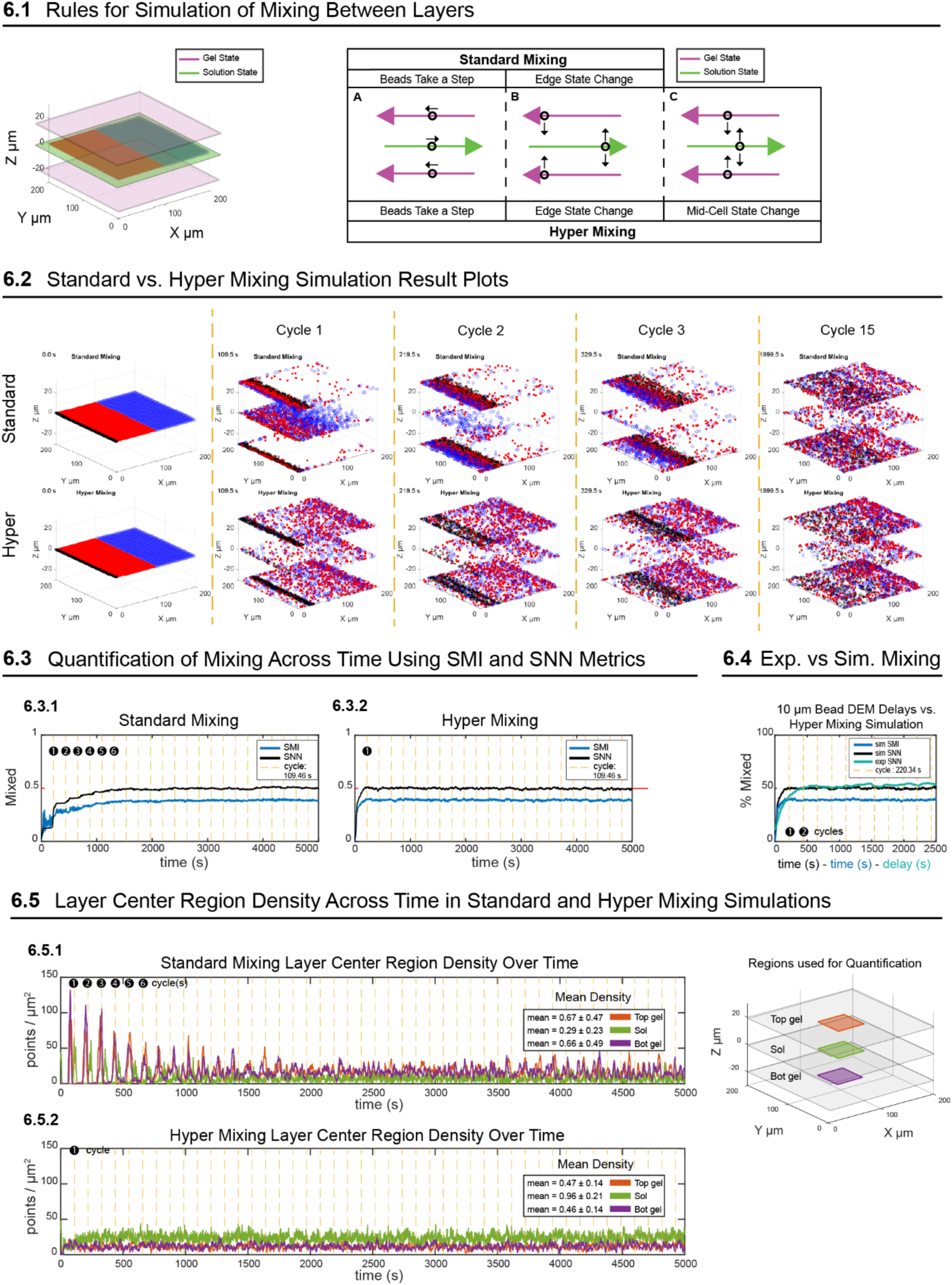
Mid-Cell Sol-Gel Transitions Accelerate Mixing. Note: Layer thickness is not represented here, and layer spacing is arbitrarily set. All transitions between layers occur in 1 jump regardless of thickness. 6.1 3D simulation model: A 3D representation of the simulation space as distinct planes, showing the gel (magenta) and liquid (green) states across the X, Y, and Z axes. 6.2 Standard vs. Hyper Mixing Simulation visual result plots. 6.3 Quantification of Mixing Across Time Using SMI and SNN Metrics: 6.3.1 SMI and SNN comparison over time for standard mixing. 6.3.2 SMI and SNN comparison over time for Hyper Mixing. 6.4 Experimental vs Simulation Mixing. SNN and SMI plotted for simulation parametrized by data in section 1. Simulation mixing completion across time plotted against delays across time for experimental data. 6.5 Layer Center Region Density Across Time in Standard and Hyper Mixing Simulations: 6.5.1 Standard mixing center point density across all 3 layers. 6.5.2 Hyper mixing center point density across all 3 layers. Hyper mixing shows a more even distribution of points across the cell layers compared to standard mixing.

The process that we simulated, referred to as hyper mixing, is governed by three core rules **(Fig. 6.1)**:

1. bead stepping within layers,
2. end switching at cell boundaries, and
3. mid-cell switching driven by dwell times.

#### Rule 1: Bead stepping (within-layer motion)

At each time step, beads are displaced according to their assigned flow state. In the sol state, bead displacement is computed independently for the x- and y-components as the sum of two terms:

1. a stochastic Brownian displacement derived from the sol diffusion coefficient, and
2. a deterministic displacement based on the mean sol velocity.

In the gel state, beads move collectively as if on a conveyor belt, taking the same deterministic step at each time point. The x-component again includes both Brownian and directed terms, but the Brownian term is derived from the gel diffusion coefficient. To reflect the confined nature of gel motion, only the y-component includes stochastic noise, and this noise is strongly reduced by scaling the gel diffusion coefficient to 1% of its measured value. This reflects the structured, cortex-associated motion of beads within the gel layer, likely driven by actin treadmilling, with the apparently diffusive term reflecting local fluctuations in treadmilling rates.

#### Rule 2: End switching (boundary transitions)

In the second rule, beads switch layer states upon reaching the cell boundaries (Y = 200 for sol flow and Y = 0 for gel flow). Upon switching states, each bead is assigned a dwell time from a dwell time distribution, and an internal counter variable is set to zero. This timer increments at successive time steps.

#### Rule 3: Mid-cell switching (dwell-time transitions)

In the third rule, beads can also switch states within the interior of the cell when their dwell time expires. Each time a bead switches state—either at the boundary or mid-cell—a new dwell time is drawn from the experimentally measured dwell time distributions to determine the next time at which it will switch states **(Figs. 5.3.5 and 5.3.6)**.

#### Control standard mixing simulation

To assess the specific contribution of mid-cell switching, we ran a simplified version of the simulation without Rule 3, a simulation we refer to as standard mixing, which includes only bead stepping and end switching **(Fig. 6.1)**. This control model corresponds to classical descriptions of sol–gel transitions in amoeba, where state changes occur primarily at the cell ends^51, 25^.

#### Mixing outcomes

We assess mixing in the simulations using two metrics: the Subdomain Mixing Index (SMI), which evaluates the fraction of different bead populations within fixed subdomains ^39^, and SNN (simple nearest neighbor, as discussed in previous sections, for experimental measurements). The SMI computes the weighted fraction of two bead populations within fixed spatial subdomains, while the SNN measures the fraction of opposite-color beads among each bead’s five nearest neighbors. The SMI is a robust metric for determining when stable mixing is achieved, as its asymptotic behavior is independent of subdomain size ^39^. We developed the SNN to provide a geometry-independent metric that does not rely on fixed subdomains, which is particularly advantageous for amoebae that continuously change shape.

In our simulations, SMI and SNN reached asymptotic values at nearly identical times, validating SNN as a reliable measure of mixing dynamics **(Figs. 6.3.1 - 6.33)**.

#### Standard vs hyper mixing performance

In the standard mixing simulation, both SMI and SNN indicated stable mixing after approximately six flow cycles **(Fig. 6.3.1)**. A flow cycle is defined as the time required for a bead to traverse the sol layer and return via the gel layer. To estimate this timescale, we initialized 1000 beads at X = 0 and measured the average time required for simulated beads to complete one full cycle (109.46 s).

Standard mixing also exhibited prolonged spatial clustering. Bead density analysis within a central subdomain showed that beads remained segregated within their original layers over extended periods **(Fig. 6.4.1)**.

In contrast, the hyper mixing simulation reached near complete mixing, indicated by the SMI and SNN, within a single flow cycle. In addition, central subdomain densities indicated uniform bead distribution after just one cycle too **(Fig. 6.4.2)** consistent with visual results **(Fig. 6.2)**.

#### Experimental validation of the simulation

The simulations above show that transient particle arrest, caused by stepping into and out of the gel layer, can drive extremely rapid “hyper-mixing.” If this model explains the behavior we observe experimentally, then simulations parameterized with values measured from living cells should reproduce the mixing timescale observed for tracer beads. To test this, we extracted dwell times, diffusion coefficients, and mean velocities directly from the same experimental video segment used to calculate the delay-based SNN mixing shown in Figure 1. We then used models fitted to these experimentally measured parameter distributions to generate simulation inputs consistent with the observed bead-tracking data, without fitting the simulation to the experimental mixing curve.

The simulated SNN curves closely matched the experimentally observed SNN curves, and simulation SMI showing complete mixing around the same time as simulation SNN and experimental SNN **(Fig. 6.3.3)**. This confirms that the simulation accurately captures the mixing dynamics observed in the amoeba cytoplasm.

Importantly, all parameter values are derived independently from bead tracking data rather than tuned to reproduce the observed mixing dynamics. This framework therefore enables quantitative investigation of intracellular mixing both indirectly through simulation and directly through experimental digital mixing analysis.

### Sol Flow Dynamics During Gel Entrance Events and Adjacent to Gel Boundary

Our measurements reveal repeated trapping and release events that, according to our simulations, can contribute to rapid intracellular mixing. One potential explanation for such apparent trapping is a shear-induced slowdown of flow near boundaries, which could transiently reduce particle velocity without requiring physical entrapment. To test whether shear-driven flow alone is sufficient to explain the observed bead dynamics, we analyzed instantaneous speed, mean speed, and mean squared displacement (MSD) for beads moving within the central sol layer of the cell, before and after they entered the gel phase.

Beads were grouped using k-means clustering based on their velocity profiles aligned to sol–gel transition points. This analysis revealed two dominant clusters **(Fig. 7.1.1)**. Cluster 1 (n = 125) consisted of an initially high-speed population characterized by a pronounced acceleration spike during the sol–gel transition **(Fig. 7.1.2)**. Rather than reflecting simple shear near a stable boundary, this transient acceleration is more consistent with a local breakdown and reorganization of the gel layer during transition. In this interpretation, weakening of the gel network may produce small, avalanche-like rearrangements of material, allowing gel material to collapse inward and adjacent material to flow in and replace it. Beads caught in this local restructuring could therefore experience a brief burst of motion as they are displaced into newly opened space or carried along with the shifting material. Cluster 2 (n = 420), in contrast, comprised a slower population with relatively steady, low velocities throughout the transition, consistent with shear-mediated slowing and possible capture near the gel boundary.

**Figure 7.**
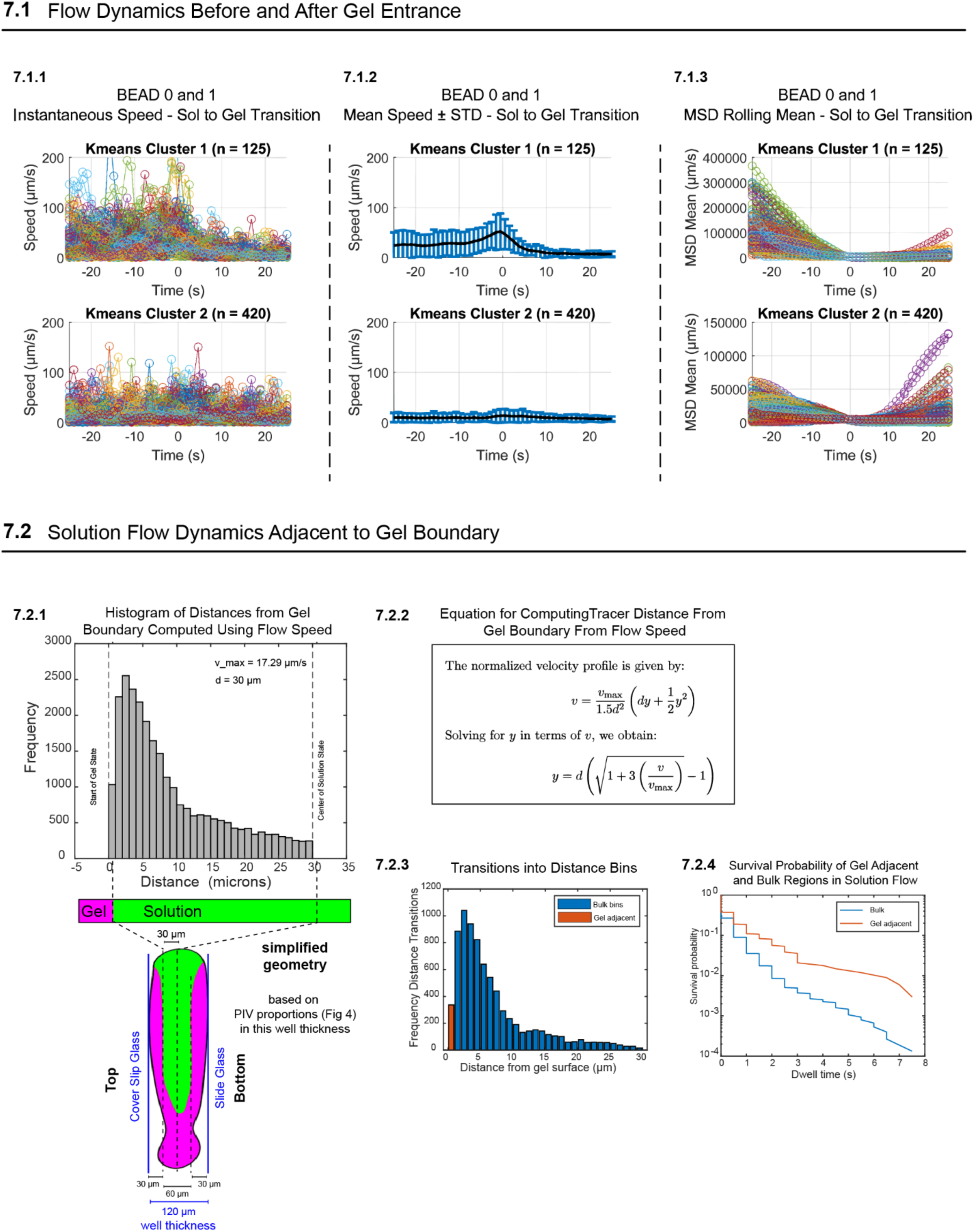
Sol Flow Dynamics During Gel Entrance Events and Adjacent to Gel Boundary. **7.1** Instantaneous bead speeds 20 seconds before and after transition from the sol to gel state. Bead 0 and Bead 1 trajectories are combined. Two populations emerge using k-means clustering (Cluster 1, high speed; Cluster 2, low speed). **7.2** Mean speed ± standard deviation for beads from Panel 7.1, aligned around the transition point. Bead 0 and Bead 1 trajectories are combined. Clusters are preserved from 7.1. **7.3** Rolling 20-second mean squared displacement (MSD) profiles of the same trajectories from 7.1, showing differences in motion type (directed vs. restricted) between clusters. **7.4** Histogram of bead distances from the gel boundary, calculated by transforming instantaneous speed values using a laminar flow model (Equation 4.2.7 from Southard, 2006) ^62^. Right panel shows the equation and its rearranged form to solve for distance y based on normalized velocity. The bottom schematic illustrates the measurement context: a central sol layer (60 µm total thickness) flanked by gel layers. The histogram reveals a concentration of beads near the gel interface, inconsistent with uniform shear flow, suggesting physical slowing or trapping at the sol-gel boundary.

To further distinguish between shear-driven slowdown and physical trapping, we computed rolling MSDs from bead trajectories aligned to sol–gel transition times **(Fig. 7.3)**. For a purely shear-driven mechanism, one would expect MSDs to decrease as beads approach the boundary and subsequently recover as they move away, with symmetric behavior of approach and separation from the surface. Instead, Cluster 1 exhibited MSD profiles indicative of sustained confinement following the transition, inconsistent with transient shear-induced deceleration. This behavior suggests that beads in Cluster 1 become physically entrapped within the gel phase following their acceleration and subsequent deceleration around the transition time (T0). By contrast, beads in Cluster 2 showed far less restricted motion after T0, consistent with a shear-driven interaction in which beads slow near the boundary but are not permanently confined and can diffuse back into the bulk sol phase.

To explicitly test whether the bead slowdown observed in Cluster 2 could be explained solely by shear-induced deceleration near the gel boundary, or whether it reflects true trapping, we used the relation between flow velocity and distance from a boundary to estimate each bead’s effective distance from the gel interface based on its instantaneous speed, allowing us to infer spatial positioning without direct distance measurements.

Using empirically determined parameters (v_m_ax = 17.29 µm/s and d = 30 µm), we converted all bead velocities into distance estimates relative to the gel surface and plotted the resulting distribution of inferred distances **(Fig. 7.2.1 and 7.2.2)**. Under pure shear-driven flow, beads would be expected to sample the sol layer uniformly as they diffuse back and forth in the sol phase in a direction perpendicular to the direction of flow. Instead, we observed a pronounced enrichment of events within 5–10 µm of the gel boundary, indicating that beads disproportionately linger near the interface. This bias toward the boundary is inconsistent with beads moving freely as implied in the shear driven transport model, and suggests the presence of additional constraint or adhesive force on bead motion near the gel surface. This behavior is illustrated schematically in figure 7.2.1, where the sol layer lies between opposing cortical gel boundaries and beads are visibly concentrated near the edges.

To determine whether the gel surface represents a dynamically distinct spatial state, we discretized the inferred bead-surface distance into equal-width bins and computed residence times corresponding to first passage transitions between neighboring bins **(Fig. 7.2.3 and 7.2.4)**. Pooling transitions from interior bins yielded a bulk reference distribution, which we compared to the distribution for the surface-adjacent bin. Beads exhibited systematically prolonged residence near the gel boundary, as evidenced by a pronounced rightward shift in the cumulative distribution function. This indicates that beads persist significantly longer in the surface-proximal region before escaping **(Fig 7.2.3 and 7.2.4)**. Such behavior is inconsistent with homogeneous diffusion or uniform advection and instead supports a model in which beads experience transient trapping or reduced mobility at the sol–gel interface, beyond what can be explained by shear alone. These results indicate that the region adjacent to the gel boundary is dynamically “sticky”: once beads enter this region, they remain there significantly longer than at other distances

### Latrunculin B Treatment Impairs Gel Dynamics and Slows Simulated Mixing

The analysis described above indicates that beads can be transiently trapped at the gel layer and then released at some later time. Actin is known to exist in different filament conformations in gel and sol layers ^30, 31^, and to be capable of undergoing sol–gel transitions. We therefore hypothesize that actin state transitions at the gel-sol interface might be involved in the transient bead trapping that contributes to intracellular mixing. To test this idea, we inhibited actin polymerization using 1 µM latrunculin B in cells injected with two 20 µm beads. Particle image velocimetry revealed a pronounced reduction in the thickness of the gel layer of Latrunculin B treated cells compared to untreated controls **(Fig. 8.1)**, confirming that actin polymerization contributes to maintaining the structural integrity and spatial extent of the gel layer. We then asked whether reduction in actin dynamics in the presence of the drug would alter the mixing dynamics of the system.

**Figure 8.**
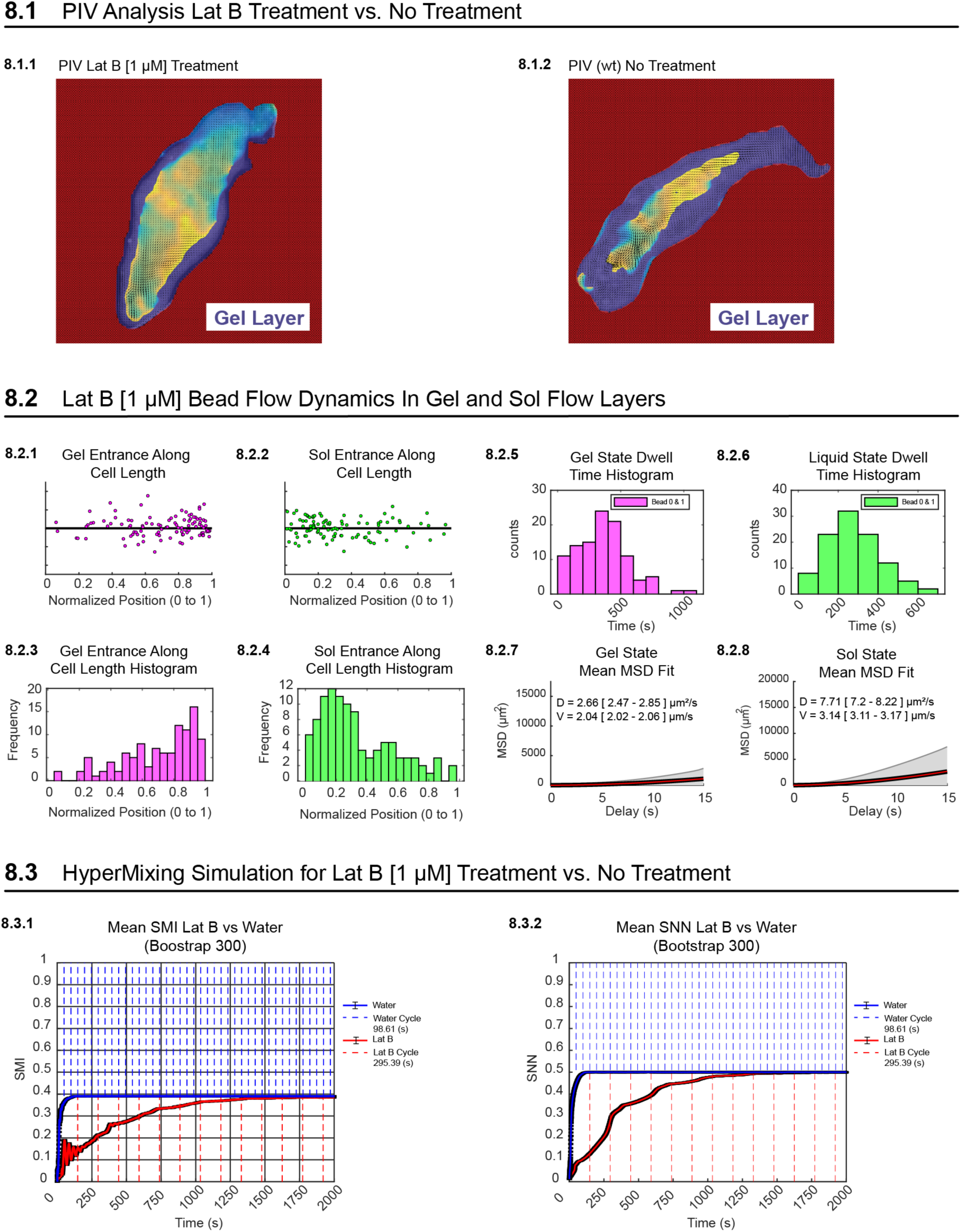
LatB Treatment Impairs Gel Dynamics and Slows Simulated Mixing. **8.1.1** PIV analysis of a cell treated with 1µM Latrunculin B. **8.1.2** PIV analysis of an untreated cell. **8.2.1- 8.2.2** Normalized layer entrance position along relative cell length plotted with jitter. **8.2.3-8.2.4** Histogram of normalized layer entrance position along relative cell length. **8.2.5-8.2.6** Histogram of bead dwell time in respective layer. **8.2.7-8.2.8** Fits of MSD used to calculate diffusion coefficient D and mean velocity V. **8.3.1** Mean SMI (Subdomain Mixing Index) across 300 simulations compared between simulations using flow parameters from latrunculin b treated cells vs untreated. **8.3.2** Mean SNN (Simple Nearest Neighbor) across 300 simulations compared between simulations using flow parameters from latrunculin b treated cells vs untreated.

As in Figure 5, we classified bead motion into gel and sol flow states to investigate layer-specific dynamics. However, Latrunculin B treatment substantially reduced the velocity contrast between the gel and sol layers, making MSD-based state classification unreliable under these conditions. To address this limitation, we developed a semi-automated classification pipeline that incorporates a spatial reference point at the uropod, allowing bead motion to be classified as retrograde or anterograde relative to the cell rear (see methods). This approach enabled robust identification of gel-associated retrograde motion and sol-associated anterograde motion even when absolute velocities in the two states were similar. Using this updated method, we measured dwell times, diffusion coefficients, and mean velocities for beads moving within each flow state.

Under Latrunculin B treatment, sol-layer entrance events were strongly enriched toward the rear of the cell **(Fig. 8.2.2 and 8.2.4)**, in contrast to untreated cells in which sol entrances are enriched toward the front **(Fig. 5.3.2 and 5.3.4)**. By comparison, gel entrance events remained distributed similarly to untreated cells, indicating that bead capture into the gel layer can still occur in the absence of robust actin polymerization. This suggests that shear-driven flow combined with residual stickiness at the gel boundary may be sufficient to remove beads from sol flow. However, unlike untreated cells, beads that entered the gel layer under Latrunculin B treatment exhibited a markedly reduced ability to return to the solution layer before reaching the rear of the cell, where transitions become forced by complete dissolving of the gel. Consistent with this interpretation, dwell time distributions increased in both gel and sol states **(Fig. 8.2.5 and 8.2.6)**, indicating slower exchange between layers.

Quantitative analysis revealed a mean velocity of 2.04 µm/s and a diffusion coefficient of 2.66 µm²/s for beads in the gel layer **(Fig. 8.2.7)**. In the sol layer, beads exhibited a higher mean velocity of 3.14 µm/s and a diffusion coefficient of 7.71 µm²/s **(Fig. 8.2.8)**. The relatively small velocity difference between layers underscores the necessity of the reference-point-based classification method, as velocity magnitude alone is insufficient to distinguish flow states under actin inhibition.

To determine whether these altered flow dynamics impact intracellular mixing, we ran our hyper-mixing simulation using layer-specific parameters measured from Latrunculin B treated cells. Simulations were repeated 300 times, and mean SMI and SNN values were compared to results from untreated simulations. In each case, 1000 additional beads incapable of mid-cell layer switching were included to determine average flow cycle duration. Latrunculin B treatment resulted in a dramatic reduction in mixing efficiency, with treated simulations requiring approximately 11 cycles to reach completion based on SMI and approximately 7 cycles based on SNN **(Fig. 8.3)**. In contrast, untreated simulations reached near-complete mixing within a single cycle and achieved complete mixing within two cycles using both metrics.

Together, these results indicate that actin dynamics are not strictly required for bead capture into the gel layer but are essential for enabling efficient return of beads to the sol layer once captured. By shifting sol entrance events toward the rear of the cell and reducing gel-to-sol exchange, actin inhibition severely impairs intracellular mixing. This supports a model in which actin-mediated dynamics sustain rapid mixing by facilitating repeated transitions between sol and gel flow states.

## DISCUSSION

In a strictly deterministic dynamical system, chaotic mixing is defined by exponential sensitivity to initial conditions, typically quantified by a positive Lyapunov exponent and arising from repeated stretching and folding of material elements. The combination of time-dependent cytoplasmic flows, repeated splitting and recombination of sol streams, and gel–sol transitions potentially could provide the necessary structural ingredients for chaotic advection.

Our measurements do not support our original model of pseudopod-driven mixing through a Baker’s transform-like mechanism. In our simulations, mixing occurred on timescales that were faster than would be expected from pseudopod formation and reabsorption alone. For example, simulations based on 20 µm bead-pair separation data predicted complete mixing in as little as 98.6 s, corresponding to approximately one flow cycle **(Fig 8.3.1).** Similarly, simulations based on bead-pair separation data from our experimental 10 µm DEM dataset predicted complete mixing in 220 s, also corresponding to approximately one flow cycle **(Fig 6.4)**. In the same 10 µm DEM dataset, complete mixing was observed experimentally within 440 s, or approximately two flow cycles **(Fig 1.4 and 6.4)**. These timescales are substantially faster than the characteristic timescale of pseudopod formation and reabsorption, which occurred over approximately 600 s in the same experimental mixing dataset.

Consistent with this interpretation, pseudopod formation was rare in the experimental mixing 10 µm DEM dataset in figure 1. Over 9.5 hours of imaging, only a single pseudopod formation and reabsorption event was observed. This suggests that this cell preferentially remained in a monopodial state, a morphology commonly associated with rapid locomotion in amoeba ^63^. Notably, the observed single pseudopod event represented an ideal case for a Baker’s transform-like mechanism: the pseudopod expanded until it contained approximately half of the cell mass before folding over being reabsorbed, as if the boundary between the folded and now touching pseudopod had merged, effectively mimicking the “split, stretch, and fold” geometry characteristic of a Baker’s transform. Despite this favorable geometry, the observed mixing dynamics were still faster than could be explained by pseudopod remodeling alone.

Moreover, when pseudopod number was experimentally increased across samples **(Fig. 3)**, bead separation rates and power-law exponents were not significantly affected. This finding suggests that pseudopod generation alone does not directly determine intracellular mixing efficiency. Thus, our results contradict the intuitive expectation that increasing pseudopod number necessarily enhances intracellular mixing. More broadly, this observation is important because several amoeboid species either rarely form pseudopods or lack prominent pseudopod-based remodeling altogether. Instead, our results suggest that efficient intracellular mixing may arise from a more general physical mechanism: the coupling of laminar cytoplasmic flows with intermittent transitions between mobile and entrapped states.

This possibility is supported by recent observations in the slime mold Physarum, another large, cytoplasmic streaming system. Although Physarum does not rely on pseudopod remodeling in the same manner as Chaos carolinensis, nuclei in Physarum have been reported to exhibit stop-and-go motion in which they intermittently become trapped within actin-based invaginations before being released back into the flowing cytoplasm ^64^. This behavior is conceptually similar to the bead dynamics observed in our study, where particles alternate between rapidly transported states and transiently immobilized or slowed states. Together, these findings suggest that intermittent entrapment and release may represent a conserved strategy for enhancing intracellular mixing in large cells with laminar cytoplasmic flows, even in the absence of pseudopod-driven folding.

In both the ballistic and super-ballistic regimes, systems exhibit high to very high mixing efficiency, and the potential for complete mixing within one cycle is strong, particularly in the super-ballistic regime where particle separation happens at an increasingly rapid rate ^41^. However, In the super-diffusive regime, while high mixing efficiency is observed, complete mixing in a single cycle is unlikely, and even more unlikely in the sub-diffusive regime. In super-diffusive regimes, particles will explore a large part of the system in a relatively short time, but the process is not as thorough as it would be in ballistic or super-ballistic regimes, where complete mixing is more probable within one cycle. Here, our data indicate that amoeba reaches near-complete global mixing within one flow cycle. From an ergodic hierarchy perspective, amoeba hyper-mixing effectively approximates Bernoulli-class mixing, the fastest class of mixing known. This observation, together with our other analyses supporting stochastic trapping and release, suggests that amoeba may have evolved a way to use laminar flows to achieve near-complete global mixing in one cycle, even though most particle pairs separate sub-ballistically. This represents one advantage that active matter mixing systems may possess over mixers made using conventional materials and strategies.

Central to our model is that the sol to gel transition occurs across the entire cell body, not just at the leading and trailing edges, as previously reported. While the mechanism controlling this transition remains unclear, our study highlights a novel characteristic: mid-cell sol to gel switching. The complete assembly and disassembly of the actin filament network at the front and rear may explain sol-gel transitions at the cell’s edges, but does not account for the newly observed mid-cell switching. Since the cortical actin network treadmills along with the gel layer from the leading to the trailing edge, we hypothesize that structural fatigue accumulates as the network travels between 1200 µm ∼ 2000 µm, depending on the shape of the amoeba, from the hyaline cap to the uropod. This fatigue could lead to breaks in the network, which are subsequently repaired through self-assembly, possibly driving mid-cell sol to gel transitions and resulting in mid-cell cytoplasmic and bead state exchanges.

Supporting this hypothesis, we found that the Birnbaum-Saunders distribution, commonly known as the fatigue life distribution, best fits the experimental gel state dwell times in experiments with larger particles like nuclei (not shown here). This distribution, typically used to model the time until failure in materials subjected to cyclic or repeated stress, suggests that the actin cytoskeleton may experience fatigue and periodic self-repair as it treadmills from front to rear. This process could explain the unique mid-cell sol to gel transitions observed and may highlight the unforeseen advantages of structural fatigue when traveling across long distances. To further investigate this possibility, future studies will aim to use live-cell fluorescence microscopy to visualize the actin cytoskeleton alongside bead dynamics, offering potential insights into this fatigue driven transition.

## METHODS

### Culturing Chaos carolinensis

*Chaos carolinensis* amoeba were cultured in 500 mL pasteurized spring water (PSW) at ambient lab temperature and light in glass containers. Amoeba were obtained from Carolina Biological Supply (Item #131324). Cultures contained *Paramecium caudatum*. (Carolina Biological Supply, Item # 131554) and rotifers introduced as contaminants of amoeba and paramecium stocks. Amoeba prey on both paramecia and rotifers. Prey populations were sustained by monthly addition of 5–10 pasteurized wheat seeds to promote bacterial growth, concurrent with replacement of 50–60% of PSW. Because amoeba localize to the bottom of the container, media exchanges were performed using a 50 mL serological pipette. Paramecium or rotifer blooms were controlled by transient addition of *Stentor coeruleus*. No predation of amoeba by *Stentor* was observed. Over time, *Stentor* were consumed by amoeba, and cultures returned to a steady-state composition of amoeba, paramecium, rotifers, wheat seeds, and bacteria.

### Microinjection

Microinjection was used to deliver 20 µm and 10 µm YG fluorescent beads (Polysciences Fluoresbrite® YG Microspheres 20.0 µm, Catalog Number 19096-2, and Polysciences Fluoresbrite® YG Carboxylate Microspheres 10.0 µm, Catalog Number 18142). All microinjections were carried out using a Drummond Nanoject II system, following our own adapted protocol for Drosophila microinjection ^35^. A Zeiss Stemi 508 stereo microscope equipped with a Transillumination 300 system and dark field illumination was utilized to visualize the microinjection needle tips easily.

To inject two 20 µm beads into amoeba, the microinjection needle was carefully cut open using a razor under a microscope until the beads could be front-filled. Once the beads were inside the needle, they were positioned near the amoeba and front-filled with low pressure to prevent them from moving too far up the needle. If the beads traveled too high, gravity guided them back down to the tip. Right before the beads were ready to fall out, the needle was swiftly inserted into the amoeba, and a minimal amount of pressure was applied to ensure precise deposition of the two beads into the amoeba’s cytoplasmic stream. The same method was employed for injecting 10 µm beads, where 18 beads were loaded and allowed to flow down the tip via gravity before being quickly injected into the amoeba to avoid premature loss of the beads.

This gravity-assisted microinjection technique was essential for the amoeba’s survival, as the large diameter of the needle needed for handling 20 µm beads carried the risk of over-injecting, which could cause the amoeba to rupture. However, using this technique, our microinjected cells closely resemble those in control amoebas injected with fluorescent dextran, which is introduced through a standard needle with an opening under 1 µm.

To further ensure amoeba health, we incubate the injected cells for one hour to allow complete wound healing, even though initial wound closure at the injection site appears to occur within seconds. Importantly, we do not anticipate any adverse effects from the size of our beads to impact the amoeba’s health.

### Microscopy

All micromanipulation was done using a Zeiss Stemi 508 with a Transillumination 300 base. This configuration is equipped with darkfield illumination which is useful for viewing microcapillary needles during micromanipulation. Injected cells were prescreened for fluorescence using a Zeis Axio Zoom V16 before mounting and imaging using other modalities. Once screened, cells were mounted in a 3 mm diameter well, created using a 120 µm thick spacer between a coverslip and a glass slide. All samples microinjected with 20 µm beads, 10 µm beads were imaged using a Nikon Ti2 Inverted Fluorescence Microscope equipped with a Nikon Plan Apo Lambda 4x Objective, a Hamamatsu ORCA-Flash4.0 v3 CMOS camera, and a quad band pass filter. We aimed to have an acquisition frame rate of 2fps while imaging in bright field and the 488 nm fluorescence channel. To achieve this frame rate, we modified our brightfield channel to fire without moving any filters from the fluorescence channel out of the way. The samples in figure 4 were imaged using an Olympus FV3000 Confocal microscope using a 60x glycerol objective.

### Image Processing

#### OttoPipeline

The OttoPipeline was developed for the automated segmentation, cropping, and registration of time-lapse videos of amoeba. This pipeline is used to turn freely crawling amoeba into amoeba crawling in place, which allows us to uncouple general cellular motion from intracellular motion (**Sup Fig 4**). The OttoPipeline consists of three separate modules, all written in MATLAB. The first module, OttoSeg, uses a maximum intensity projection to generate a background image, which is then subtracted from every frame in the video to produce segmented frames. Variations of this approach include using a median intensity projection to compensate for fluctuations in transmitted light sources, and a rolling projection to generate a series of background images over time. The rolling projection is especially useful for cases where bubbles form inside the spacer during long-lapse acquisitions, helping to maintain accurate segmentation. The second module, OttoCrop, takes the segmented frames from OttoSeg and binarizes each one to generate a centroid, which is then used for automated cropping. The user defines the crop height and width, which are applied uniformly to all frames. In the final module, OttoReg, performs image registration across time to refine alignment using a configuration created with MATLAB’s Image Registration Estimator tool.

All modules have been optimized for speed to handle the 10 TB of data generated from our long timelapse image acquisitions. To achieve this, the pipeline is designed to support multicore processing using MATLAB’s Parallel Computing Toolbox.

### Computational Analysis

#### Geodesic distance

Our analysis of mixing dynamics is based in part on pairwise distances between beads, in which the temporal dynamics with which such distance increases can be used to classify the type of fluid motion. Using geodesic distance to measure distances within our amoeba’s cytoplasm is essential for accurately capturing spatial relationships as the cell is not convex and constantly changes shape and extends multiple pseudopods. Unlike Euclidean distance, which measures a straight-line path, geodesic distance follows the actual curved route along the amoeba’s body, reflecting the shortest path through its complex, shifting, and non-convex landscape. This distinction is illustrated in **Supplementary Figure 5** where the Euclidean distance between two points in the amoeba (left panels) differs significantly from the geodesic distance (right panels). The Euclidean measure, shown as a direct line, fails to account for the amoeba’s contours and structural boundaries. In contrast, the geodesic distance follows the organism’s curved shape, adapting as the amoeba changes form. This approach provides a more realistic and meaningful measurement of separation within the dynamic environment of the amoeba, accommodating its irregular morphology and continuous structural changes. While more accurate, we note that the geodesic distance calculation takes significantly longer than the Euclidean distance calculation. This is because the distance between the bead and every possible pixel must be calculated for each frame.

#### Continuous bead tracking

Bead tracking was performed using the TrackMate plugin in ImageJ ^36^, with subsequent analysis carried out in MATLAB by loading the spot and track data into spreadsheets. Tracking multiple particles presents a significant challenge, especially when more than two beads are microinjected into a cell, as it complicates continuous trackability. For example, tracking two beads can produce two trajectories that span the entire video. However, when tracking multiple beads, accuracy decreases, often causing tracks to break mid-trajectory. This results in multiple tracks being generated for a single bead.

In our dataset, which contained 18 microinjected 10 µm beads, we generated a total of 54 tracks. To analyze this video effectively, we developed a MATLAB script that identifies overlapping tracks. This allowed us to isolate video segments containing 18 continuous tracks that span the entire length of the video segment. Using this method, we were able to digitally label two distinct bead populations based on their known trajectories, enabling us to study how quickly these two labeled populations mixed over time.

This algorithm allowed us to observe multiple mixing events within a single video and gave us the flexibility to analyze mixing starting at any chosen frame. We leveraged this to examine mixing across different delay times.

#### Layer Classification via Rolling MSD Threshold

To study mixing from two bead trajectories, we track them over time and extract key parameters such as the diffusion coefficient, mean velocity, and dwell time for the respective gel and liquid states. To classify these states, we first apply a 20-second rolling mean squared displacement (MSD) across each bead’s trajectory, generating an array where each frame corresponds to the MSD across time within the 20 second window. We then apply a threshold against the 20 second window MSD to distinguish gel states from liquid states. The accuracy of this threshold is verified post-analysis by generating a video overlay that highlights the gel and sol/liquid state trajectories.

Once the gel and liquid segments are identified for each trajectory, MSD analysis is applied to these track subsets to derive diffusion coefficients and mean velocities by fitting a diffusion with drift model to the MSD curve for each state. These parameters are then used to simulate bead movement, allowing us to assess mixing based on the simulation results.

#### Layer Classification via Reference Point

To distinguish anterograde sol phase flow from retrograde gel flow, a reference point corresponding to the trailing edge of the cell was defined at each time frame. This reference was computed by fitting a line through a sequence of trailing centroid positions and identifying the intersection of this line with the segmented cell boundary. During periods of near-linear cell motion, the centroid-derived line reliably intersected the posterior edge of the cell, providing a stable trailing-edge reference. During sharp cell turns, however, the centroid line shifted laterally and no longer intersected the true posterior boundary. These mislocalized reference points were corrected semi-automatically by manually selecting corrected trailing-edge positions at 50-frame intervals within the affected time windows and interpolating the trailing-edge position across the intervening frames.

To classify bead motion relative to the trailing-edge reference, we computed the bead–reference distance time series for each bead,

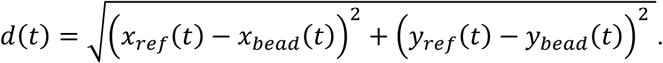

Distance traces were smoothed using a locally weighted regression (LOWESS) filter. The frame-to-frame change in the smoothed distance, was used as a signed proxy for motion along the anteroposterior axis: increasing distance (*v*(*t*) > *ε*) was classified as sol/anterograde flow (+1), decreasing distance (*v*(*t*) < −*ε*) as

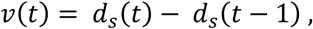

gel/retrograde flow (−1), and small-magnitude changes (∣ *v*(*t*) ∣≤ *ε*) were treated as neutral (0). A deadband threshold of ε=0.05 μm/frame was applied to suppress noise-driven sign changes. To remove short spurious bursts, sol and gel classifications were additionally filtered by enforcing a minimum run length of 50 frames, after which neutral frames were imputed by carrying forward the most recent non-neutral state to yield a continuous flow-state trace for each bead.

#### Baker’s Transformation Simulation and Power Law Analysis

To provide a theoretical baseline for cytoplasmic mixing, we simulated bead advection under a discrete Baker’s Transformation, implemented in a two-dimensional 200×200 μm domain. An initial lattice was populated with *N*=10000 beads, randomly sampled to fill the square domain. Small spatial perturbations were introduced by adding uniform noise of amplitude *A* = 2 μm to both *x* and *y* coordinates to avoid artificial symmetry and clumping.

Each cycle of the Baker’s Transformation consisted of a stretch step followed by a fold step. In the stretch step, positions were updated as:

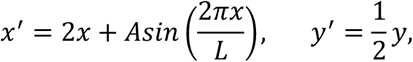

where *L*=200 μm is the domain size. The sinusoidal term provides a small perturbation to mimic biological irregularities and prevent line-based clustering.

In the fold step, beads crossing the right boundary were shifted back into the domain and displaced upward:

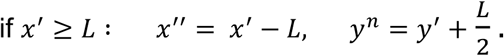

After each fold, coordinates were clamped and linearly rescaled so that all bead positions spanned the interval [0, *L*] in both *x* and *y*. This normalization prevents drift and ensures that bead density remains uniform across cycles.

Cycle boundaries were defined immediately after the fold step. Simulations were run for 100 cycles, corresponding to a total time of 12000 s (120 s per cycle). Bead positions were saved at t=0,120,…,12000 s.

Nearest neighbors (NN) were assigned once at *t*=0 by computing the Euclidean distance matrix among all bead positions and selecting the closest non-self neighbor for each bead. For

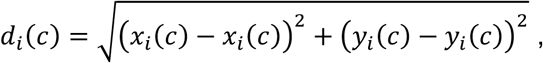

bead *i* with neighbor *j*, the pairwise separation at cycle *c* was defined as: where (*x_i_*(*c*), *y_i_*(*c*)) denotes the position of bead I at cycle *c*.

For each bead pair, the temporal evolution of separation was tested for a power law relationship of the form:

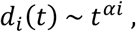

where *t* is time (in seconds) and *αi* is the fitted exponent describing the rate of separation. To estimate *αi*, we performed linear regression of the log–log transformed relationship across all cycle boundaries:

using *t* = 120, 240,…,12000 s. Bead pairs with nonpositive separations, negligible variance (std(*d_i_*)<0.01), or extreme slopes (∣*α_i_*∣>10) were excluded.

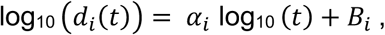

The set of valid exponents *α_i_* was analyzed by kernel density estimation (KDE) to obtain the probability density distribution. Mean and median exponents were reported as summary statistics. In the baseline configuration, 9949 of 10000 bead pairs yielded valid fits, producing a unimodal distribution centered at *α* ≈ 0.5.

#### Quantifying mixing SMI and SNN metrics

To quantitatively measure mixing across time in our simulation data and experimental data, we use two distinct mixing metrics. The first, the Subdomain Mixing Index (SMI), evaluates the fraction of different bead populations within fixed subdomains ^39^. Subdomain-based methods divide the system into smaller, defined regions or “bins” to quantify the distribution of distinct bead populations within each subdomain. This approach evaluates mixing by comparing the relative proportions of bead types in each region to an ideal mixed state. In the SMI, each subdomain is weighted based on the number of beads it contains relative to the total population. The sum of these subdomain fractions gives the SMI. Although the SMI was formulated to handle multiple distinct populations within a given space^39^, we have simplified the SMI formula to handle two distinct populations.

The SMI metric is particularly useful for measuring mixing in fixed spaces, like those represented in our simulations. The SMI accurately determines the length of time it takes for a system to reach a stable mixing state ^39^. While adjusting subdomain size can affect the overall mixing magnitude reported, it has no impact on the time it takes to reach a stable state. Under the assumption that particles in two different marked populations are initially present in equal numbers, the SMI would start close to zero and then approach 0.5 as mixing reaches its asymptotic level.

The SMI is particularly useful for measuring mixing in fixed spaces, such as those represented in our simulations, and it accurately captures the time required for a system to reach a stable mixing state.^39^ While adjusting subdomain size can affect the overall mixing magnitude reported, it has no impact on the time it takes to reach a stable state. Under the assumption that particles in two different marked populations are initially present in equal numbers, the SMI would start close to zero and then approach 0.5 as mixing reaches its asymptotic level. Unlike Cho et al. ^39^, we do not normalize this value to 1. Such scaling is useful in multicomponent systems, where the raw maximum depends on the number of species being compared. Because our analysis tracks only two populations, the unscaled maximum of 0.5 is the correct upper bound and provides a more direct interpretation of the binary mixing state.

Our second metric, the Simple Nearest Neighbor (SNN) score is designed for dynamic environments, such as inside a shape-changing amoeba. Neighbor-based methods to calculate mixing focus on the immediate surroundings of individual beads, analyzing how the local environment contributes to overall mixing. The SNN metric calculates the fraction of mixing by summing the fractions of similarly labeled beads from the nearest five neighbors for each bead, thereby avoiding the limitations of fixed space, making this metric more directly applicable to live-cell data. While SNN ranges from 0 to 1, a value of 0.5 indicates random mixing of two equally represented populations, whereas values approaching 1 indicate increasing local order, with 1 corresponding to a perfect checkerboard pattern. This makes comparison with SMI straightforward, since for a two-population system both metrics span the 0 to 0.5 range when describing the progression from an initially separated state to random mixing. Formulas for our SMI and SNN metric can be found below:

<u>Formula 1: Subdomain Mixing Index (SMI)</u>

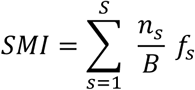

Let:

- *S* = 192 be the total number of subdomains. (64 subdomains per layer)
- *B* be the total number of beads.
- *n_s_* be the number of beads in subdomain *s*, where *s* = 1,2,…, *S*.
- *f_s_* be the smaller fraction of the two different bead populations within subdomain *s*.
- Each subdomain is weighted by the ratio 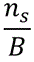

Where:

- 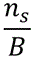is the weight of subdomain *s*, representing its proportion relative to the total population.
- *f_s_* is the smaller fraction of the two different bead populations within subdomain *s*.

In other words, the SMI is calculated by summing the weighted fractions of different bead populations across all 192 domains.

<u>Formula 2: Simple Nearest Neighbor (SNN)</u>

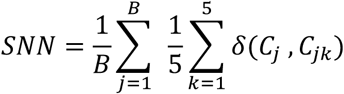

Let:

- *B* be the total number of beads.
- For each bead *j* (where *j* = 1,2, …, *B*), we find the five nearest neighbors using Euclidean distance
- For each of the 5 nearest neighbors of bead *j*, we create bead pairs and score them as follows:

- If a bead pair has the same color, it scores 0.

- If a bead pair has different color, it scores 1.

Where:

- *Cj* is the color of bead *j*.
- *Cjk* is the color of the *k*-th nearest neighbor of bead *j*
- *δ*(*C*_j_, *C* _j*k*_) is an indicator function that outputs 1 if *C_J_* ≠ *C*_j*k*_ (different colors) and 0 if *C_J_* = *C*_j*k*_ (same color).

### Hyper mixing simulation

We used a structured element matrix in MATLAB to model the movement of 2,500 beads, divided into two groups of 1,250. At T0, all beads are evenly distributed over a 200 µm x 200 µm plane at z = 0, representing the sol layer. This layer is split across the middle of the Y axis to initiate blue and red groups at T0. As time progresses, this sheet moves in the positive X direction, with additional layers at z = 20 and z = −20 representing the exterior gel layers of the amoeba. For each bead, its XY position and layer (upper gel, lower gel, or sol associated) is stored, along with a clock variable that tracks the time until the next state transition between layers. In addition to the switching clock, beads switch layers when they flow to the end of their layer. All layer switches assign new dwell times.

The simulation is carried out in three primary sections, each implementing specific rules. The first section, the “stepping section,” moves all beads in their respective directions; sol beads flow in the positive X direction, while gel beads flow in the negative X direction as the gel treadmills backwards in the reference frame of the cell. Here movement in the Y direction is driven through diffusion to reflect the properties of laminar flow. We used a Gaussian model based on Einstein’s formula for diffusion to simulate these steps ^37,38^. The step size for each bead is determined using a random number drawn from a Gaussian distribution, fitted to the experimentally derived diffusion coefficient and mean velocity ranges.

In the second section, we check for beads that have reached the end of their layers. Beads reaching the end of their layer will switch states, moving from sol to gel or vice versa. In our simulations, the probability of switching to either gel layer is set to 50% since we don’t have experimental data to fit a distribution to model the gel layer selection.

In the third section, beads that have switched states are assigned a clock state drawn from a dwell time distribution fit to our experimental data. For all beads, the clock is decremented each time-step, and when the clock reaches zero, the bead switches states regardless of whether they have reached the end of their current layer. Once this happens a new value is updated, and the clock is reset. The first clock value is assigned for every bead at T0. Clock values are generated using a random number, fitted to a distribution based on experimental dwell time data for gel and sol layers. The most appropriate distribution is selected by testing all distributions available in MATLAB’s function MLE (including: Weibull, Nakagami, gamma, gp, gev, birnbaumsaunders, Rayleigh, rician, lognormal, loglogistic, inverse gaussian, uniform, normal, tlocationscale, logistic, exponential) and choosing the one with the lowest AIC (Akaike information criterion) value which optimizes fit taking into account the number of free fitting parameters, which differs among distribution forms.

To assess the impact of layer switching caused by the third section of our simulation, we ran the simulation without this feature, referring to it as “standard mixing.” When all three sections are enabled, including the layer switching, we refer to it as “hyper mixing.”

## ACKNOWLEDGMENTS

We thank the members of the Marshall Lab, the 2023 Physiology course at MBL Woods Hole, and the 2024 CCC Summer Course at San Francisco State University, for meaningful discussions and contributions towards this project. We further thank Greyson Lewis and Nat Handle from the Marshall lab for insights into image processing and computational modeling, as well as Harry Tuazon and Lakshmi Balasubramaniam, from the 2023 MBL physiology course, for meaningful discussions on particle image velocimetry and the application of Eulerian vs Lagrangian approaches. We also thank students from the CCC Summer Course for their work on nuclei mixing dynamics (data not shown here). This work was supported by the HHMI Gilliam program (UD), and NIH grant R35 GM130327 (WFM). Work done by students in the CCC Summer Course was supported by the Center for Cellular Construction funded by NSF grant DBI-1548297.

## SUPPLEMENTARY FIGURES

**Supplementary Figure 1.**
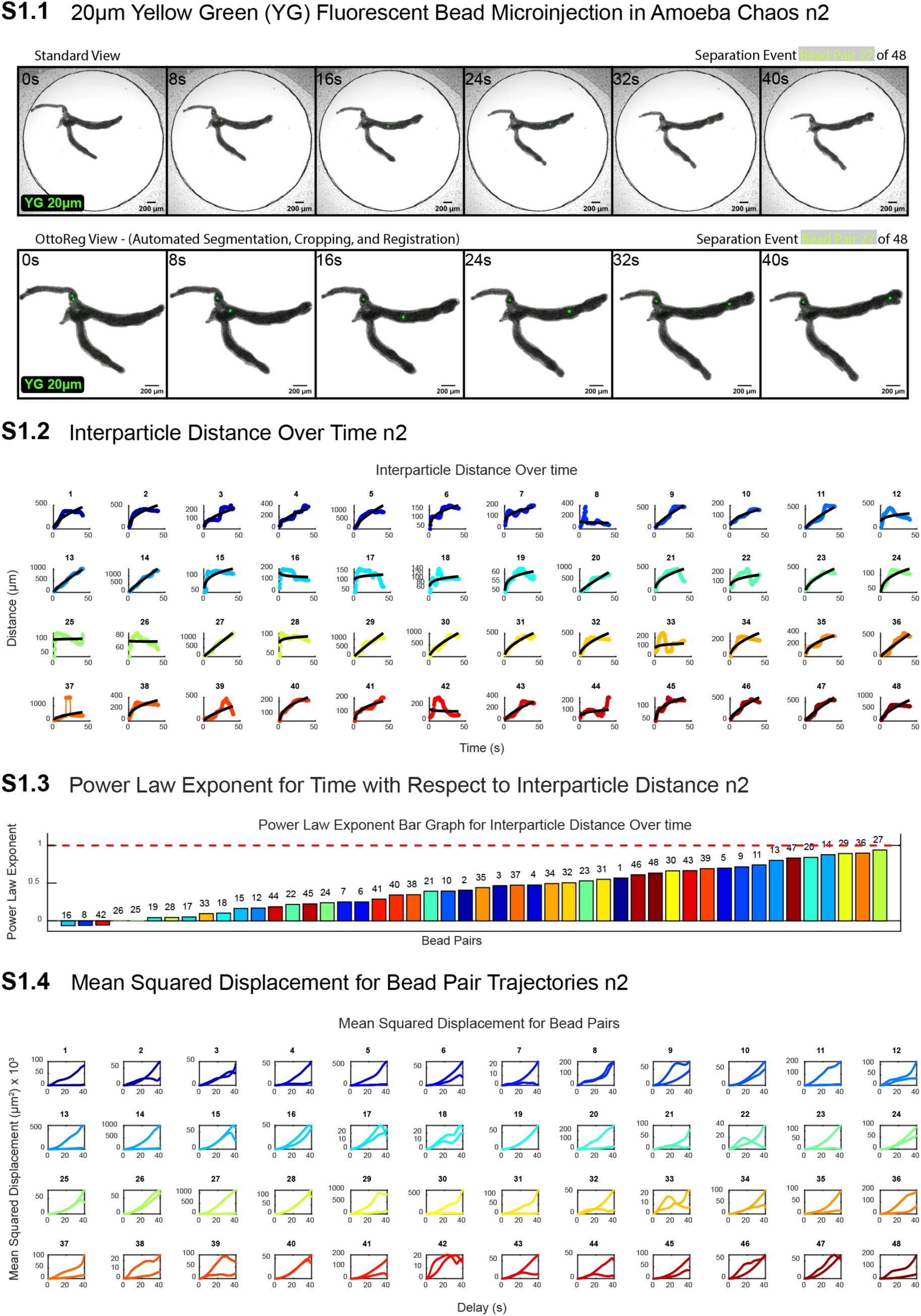
20 µm Bead Separation Assay n2 of 4 S1.1 Time-lapse images showing the microinjection of 20 µm yellow-green (YG) fluorescent beads into amoeba *Chaos*. The top row presents the standard view, and the bottom row shows the OttoReg view, where automated segmentation, cropping, and registration have been applied. The images capture a separation event for bead pair 21, showing the displacement of the beads over a 40-second time window. Scale bars represent 200 µm. S1.2 Interparticle distance over time for bead pairs tracked in the experiment. Each plot corresponds to a unique bead pair, displaying the distance (µm) between beads over time (s). S1.3 Power law exponent for interparticle distance as a function of time. The bar graph shows the power law exponent for each bead pair, with values color-coded and labeled. The red dashed line at Y = 1. S1.4 (MSD) plots for each bead pair trajectory. These plots show the MSD (µm²) as a function of delay time (seconds), providing insight into the diffusion behavior of the beads within the amoeba.

**Supplementary Figure 2.**
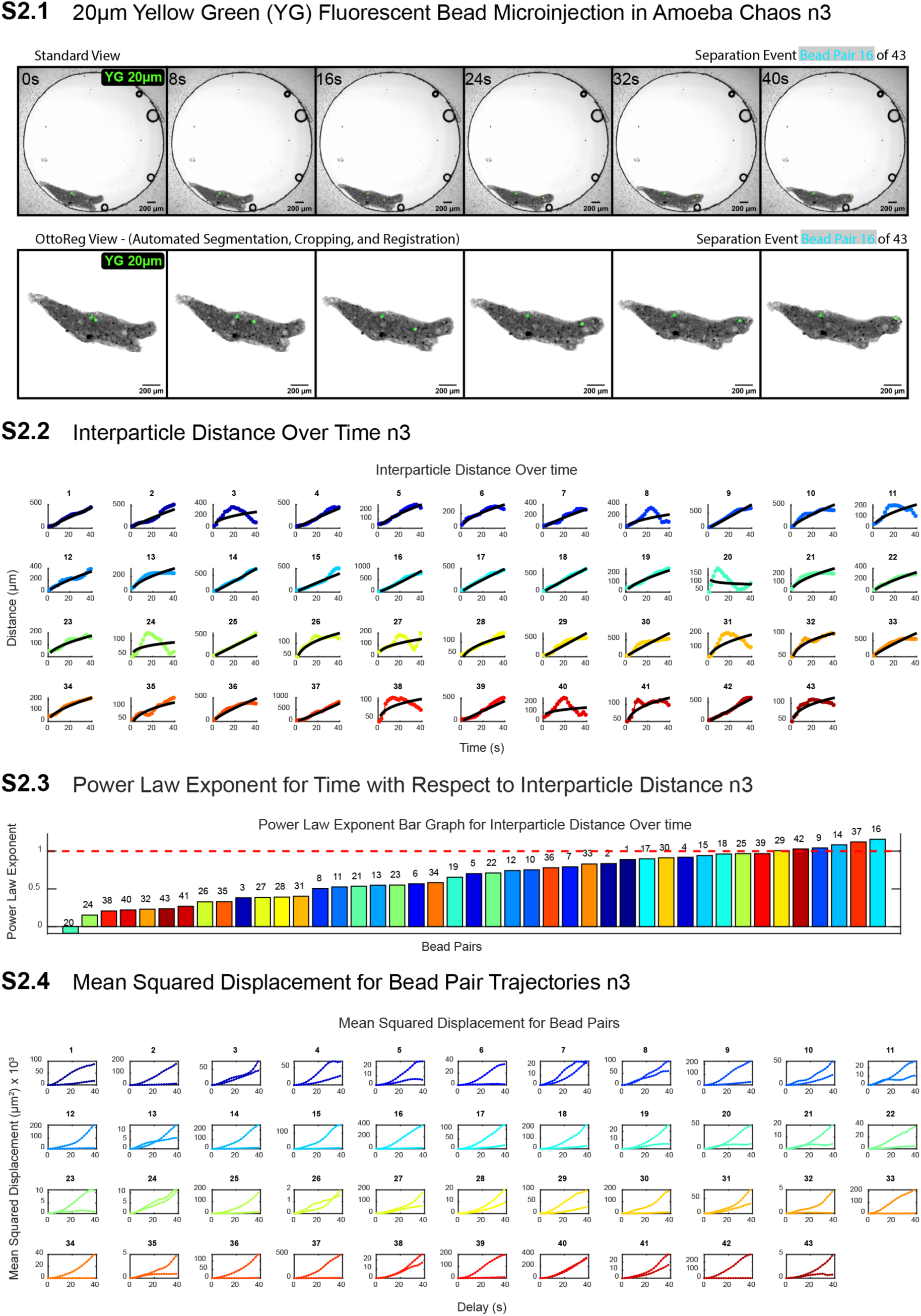
20 µm Bead Separation Assay n3 of 4 S2.1 Time-lapse images showing the microinjection of 20 µm yellow-green (YG) fluorescent beads into amoeba *Chaos*. The top row presents the standard view, and the bottom row shows the OttoReg view, where automated segmentation, cropping, and registration have been applied. The images capture a separation event for bead pair 16, showing the displacement of the beads over a 40-second time window. Scale bars represent 200 µm. S2.2 Interparticle distance over time for bead pairs tracked in the experiment. Each plot corresponds to a unique bead pair, displaying the distance (µm) between beads over time (s). S2.3 Power law exponent for interparticle distance as a function of time. The bar graph shows the power law exponent for each bead pair, with values color-coded and labeled. The red dashed line at Y = 1. S2.4 (MSD) plots for each bead pair trajectory. These plots show the MSD (µm²) as a function of delay time (seconds), providing insight into the diffusion behavior of the beads within the amoeba.

**Supplementary Figure 3.**
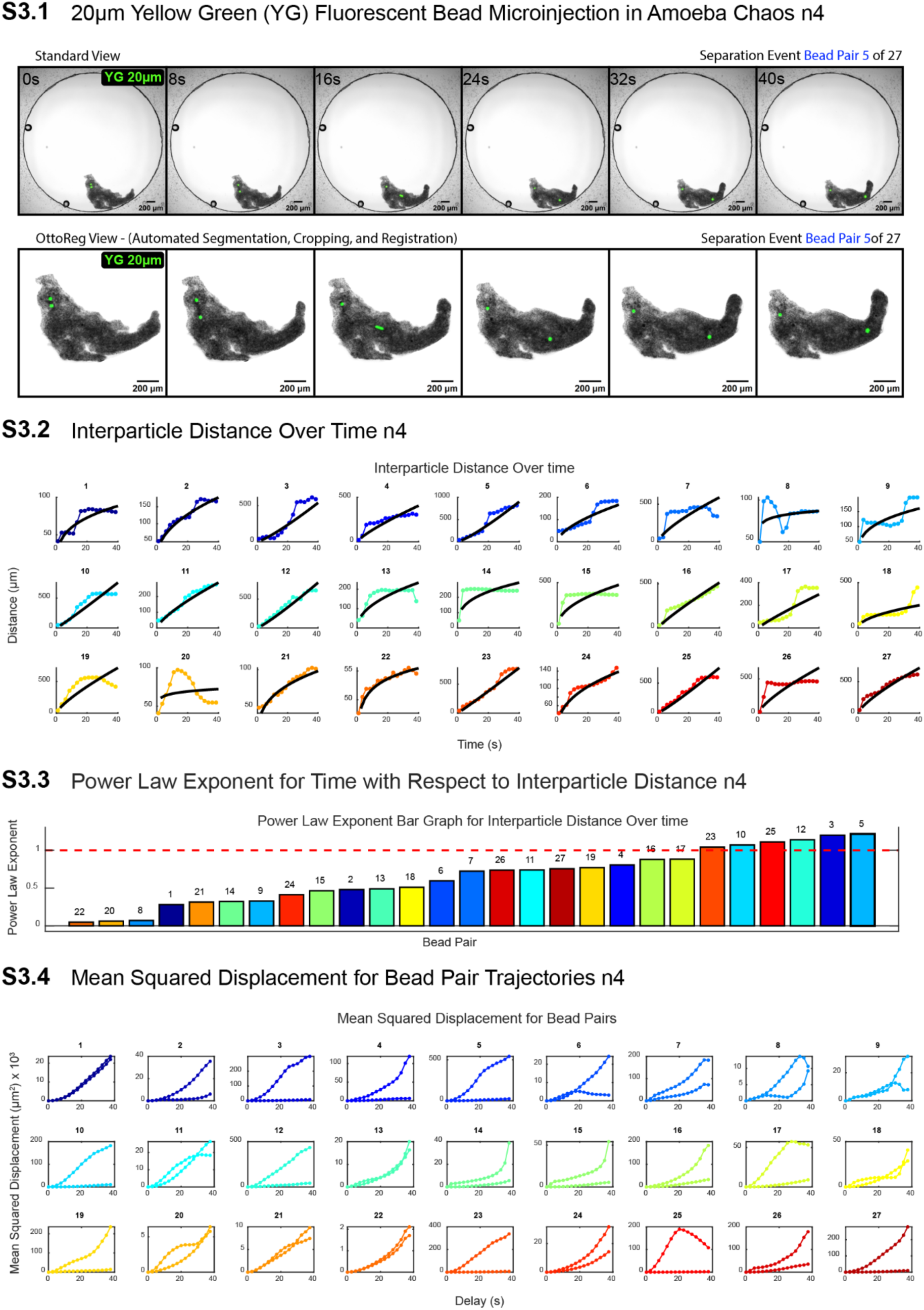
20 µm Bead Separation Assay n4 of 4 S3.1 Time-lapse images showing the microinjection of 20 µm yellow-green (YG) fluorescent beads into amoeba *Chaos*. The top row presents the standard view, and the bottom row shows the OttoReg view, where automated segmentation, cropping, and registration have been applied. The images capture a separation event for bead pair 5, showing the displacement of the beads over a 40-second time window. Scale bars represent 200 µm. S3.2 Interparticle distance over time for bead pairs tracked in the experiment. Each plot corresponds to a unique bead pair, displaying the distance (µm) between beads over time (s). S3.3 Power law exponent for interparticle distance as a function of time. The bar graph shows the power law exponent for each bead pair, with values color-coded and labeled. The red dashed line at Y = 1. S3.4 (MSD) plots for each bead pair trajectory. These plots show the MSD (µm²) as a function of delay time (seconds), providing insight into the diffusion behavior of the beads within the amoeba.

**Supplementary Figure 4.**
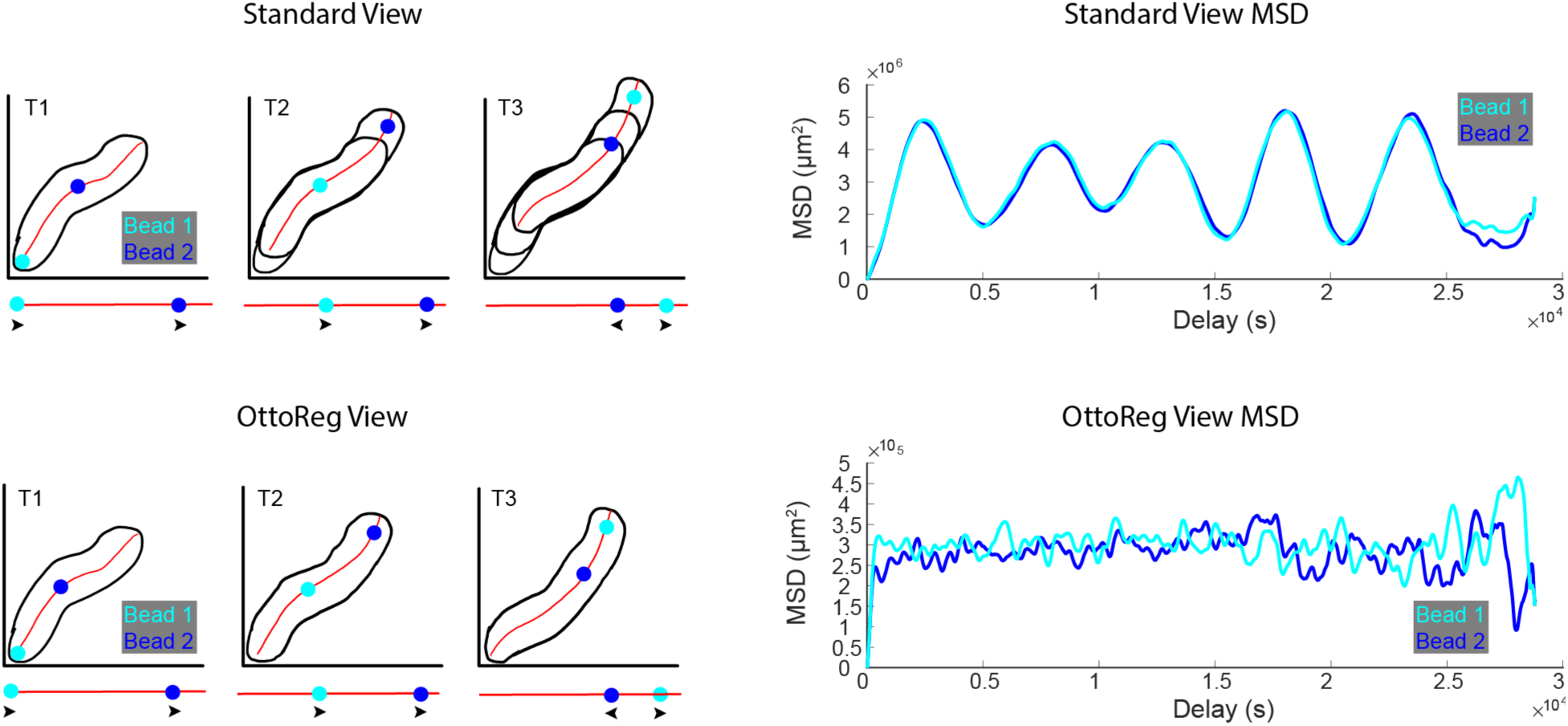
Separating Bead Movement from Cell Movement Using OttoReg This figure demonstrates the effect of the OttoReg pipeline on separating bead movement from cell movement during tracking. The left panels (T1, T2, T3) show the trajectories of two tracked beads (cyan for Bead 1 and blue for Bead 2) over time in both the Standard View (top row) and the OttoReg View (bottom row). In the Standard View, both bead movement and overall cell movement are conflated, as shown by the shifting cell outline and bead positions. The OttoReg View corrects for cell movement, isolating the true motion of the beads within the cell. The right panels show the mean squared displacement (MSD) plots for both beads in the Standard View and OttoReg View. In the Standard View MSD, oscillations are evident, likely due to cell movement, while in the OttoReg View, these oscillations are reduced, providing a clearer representation of the actual bead motion independent of the cell’s overall movement.

**Supplementary Figure 5.**
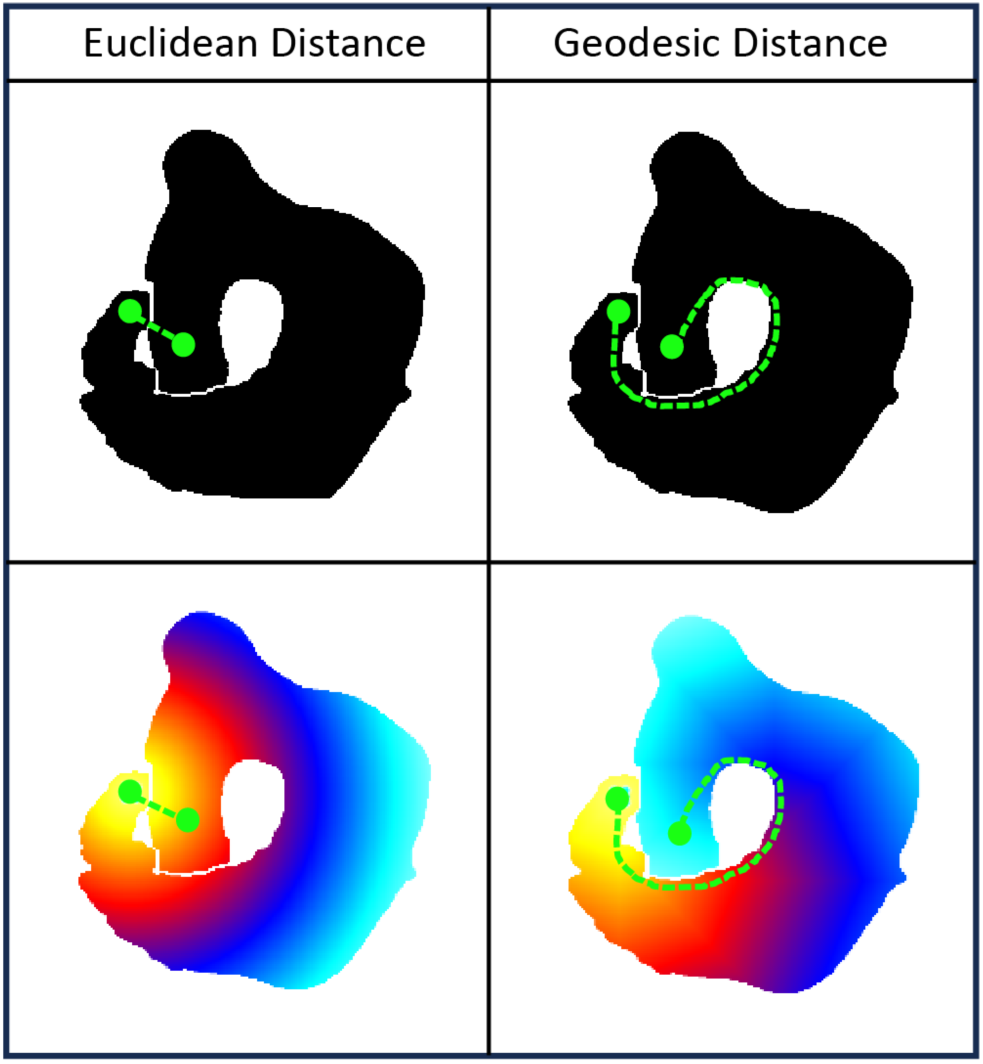
Euclidean Distance vs. Geodesic Distance This figure compares the differences between Euclidean distance and geodesic distance within a binary image of an amoeba. The left column illustrates the Euclidean distance, where the shortest path is calculated in a straight line between two points, even if obstacles are present. The right column shows the geodesic distance, which accounts for the object’s geometry and calculates the shortest path along the surface. The top row displays the binary representation of the two-distance metrics, while the bottom row shows the corresponding distance gradients with color-coded heat maps, where warmer colors represent shorter distances, and cooler colors represent longer distances. The geodesic distance, as shown, correctly follows the contour of the shape, while the Euclidean distance disregards the object’s boundaries.

## Notes

### Competing Interest Statement

The authors have declared no competing interest.

